# Whole-genome and RNA sequencing reveal variation and transcriptomic coordination in the developing human prefrontal cortex

**DOI:** 10.1101/585430

**Authors:** Donna M. Werling, Sirisha Pochareddy, Jinmyung Choi, Joon-Yong An, Brooke Sheppard, Minshi Peng, Zhen Li, Claudia Dastmalchi, Gabriel Santpere, Andre M. M. Sousa, Andrew T. N. Tebbenkamp, Navjot Kaur, Forrest O. Gulden, Michael S. Breen, Lindsay Liang, Michael C. Gilson, Xuefang Zhao, Shan Dong, Lambertus Klei, A. Ercument Cicek, Joseph D. Buxbaum, Homa Adle-Biassette, Jean-Leon Thomas, Kimberly A. Aldinger, Diana R. O’Day, Ian A. Glass, Noah A. Zaitlen, Michael E. Talkowski, Kathryn Roeder, Matthew W. State, Bernie Devlin, Stephan J. Sanders, Nenad Sestan

## Abstract

Variation in gene expression underlies neurotypical development, while genomic variants contribute to neuropsychiatric disorders. BrainVar is a unique resource of paired whole-genome sequencing and bulk-tissue RNA-sequencing from the human dorsolateral prefrontal cortex of 176 neurotypical individuals across prenatal and postnatal development, providing the opportunity to assay genomic and transcriptomic variation in tandem. Leveraging this resource, we identified rare premature stop codons with commensurate reduced and allele-specific expression of corresponding genes, and common variants that alter gene expression (expression quantitative trait loci, eQTLs). Categorizing eQTLs by prenatal and postnatal effect, genes affected by temporally-specific eQTLs, compared to constitutive eQTLs, are enriched for haploinsufficiency, protein-protein interactions, and neuropsychiatric disorder risk loci. Expression levels of over 12,000 genes rise or fall in a concerted late-fetal transition, with the transitional genes enriched for cell type specific genes and neuropsychiatric disorder loci, underscoring the importance of cataloguing developmental trajectories in understanding cortical physiology and pathology.

**Highlights:**

- Whole-genome and RNA-sequencing across human prefrontal cortex development in BrainVar
- Gene-specific developmental trajectories characterize the late-fetal transition
- Identification of constitutive, prenatal-specific, postnatal-specific, and rare eQTLs
- Integrated analysis reveals genetic and developmental influences on CNS traits and disorders

## Introduction

The human nervous system develops slowly over several decades, starting during embryogenesis and extending postnatally through infancy, childhood, adolescence and young adulthood (Keshavan et al., 2014; Shaw et al., 2010; Silbereis et al., 2016; Tau and Peterson, 2010). Over this time, myriads of functionally distinct cell types, circuits, and regions are formed (Hu et al., 2014; Lui et al., 2011; Silbereis et al., 2016). These aspects of neurodevelopment are highly context-dependent, since neural cells are born in an immature state and must differentiate, migrate, and establish circuits to produce regionally distinct brain structures. In doing so, the cells undergo a variety of molecular and morphological changes. Consequently, the characteristics of a given cell and brain region at a given time offer only a snapshot of organogenesis and brain function, necessitating consistent profiling across development.

The molecular and cellular processes underlying development of the nervous system rely on the diversity of transcripts and their precise spatiotemporal regulation (Bae and Walsh, 2013; Silbereis et al., 2016). Functional genomic analyses of the developing human brain have revealed highly dynamic spatiotemporal patterns of gene expression and epigenetic changes during prenatal and early postnatal development (Kang et al., 2011; Li et al., 2018). In contrast, these metrics are comparatively stable over several decades of adulthood (Colantuoni et al., 2011; Jaffe et al., 2018; Kang et al., 2011; Li et al., 2018; Pletikos et al., 2014). Disruptions of these developmentally dynamic regulatory processes are likely to be important contributors to multiple neurodevelopmental and neuropsychiatric disorders (Birnbaum and Weinberger, 2017; Breen et al., 2016; Geschwind and Flint, 2015; McCarroll and Hyman, 2013; Rosti et al., 2014; Sestan and State, 2018; Turner and Eichler, 2018). In keeping with this expectation, analyses of spatiotemporal expression patterns have implicated mid-fetal brain development as a vulnerable process, and the prefrontal cortex as a vulnerable region, for both autism spectrum disorder (ASD) and schizophrenia (SCZ) risk genes (Chang et al., 2015b; Gulsuner et al., 2013; Li et al., 2018; Network and Pathway Analysis Subgroup of the Psychiatric Genomics Consortium, 2015; Parikshak et al., 2013; Satterstrom et al., 2018; Willsey et al., 2013; Xu et al., 2014). More generally, atypical trajectories of brain maturation likely affecting pathophysiological manifestations have been described in ASD, SCZ, and other neuropsychiatric disorders (Birnbaum and Weinberger, 2017; Courchesne et al., 2007; Ecker et al., 2015; Insel, 2010; Keshavan et al., 2014; Shaw et al., 2010; Tang and Gur, 2018). Given that neuropsychiatric disorders have discrete ages of onset and patterns of progression and may arise due to genetic or environmental insults at various times and places during the life of an individual, there is a clear need to examine gene expression and neuropsychiatric risk across the span of human brain development.

In addition to spatiotemporal variation, genetic sequence variants are also associated with variation in gene expression levels, which can thereby contribute to differences in brain structure, function, and behavior between individuals (Elliott et al., 2018). Several laboratories and consortia have systematically identified such expression quantitative trait loci (eQTLs) through paired analysis of single nucleotide polymorphism (SNP) genotyping and gene expression data in numerous tissues, including the brain (BrainSeq: A. Human Brain Genomics Consortium, 2015; Dobbyn et al., 2018; Fromer et al., 2016; Gibbs et al., 2010; Heinzen et al., 2008; Jaffe et al., 2018; Liu et al., 2010; Myers et al., 2007; Psych et al., 2015; The GTEx Consortium, 2015; Wang et al., 2018). However, there are relatively few such studies of the developing human brain (Colantuoni et al., 2011; Jaffe et al., 2018; Kang et al., 2011; O’Brien et al., 2018) and most of these, as a result of limitations forced by then-contemporary technologies or limited tissue availability, were unable to assess the full scope of genetic variants and/or gene expression dynamics across the whole of brain development (Colantuoni et al., 2011; Jaffe et al., 2018; Li et al., 2018; O’Brien et al., 2018). Thus, developmentally regulated and rare variant eQTLs are sparsely represented in the current catalog of human brain eQTLs, highlighting the need for additional resources. Such eQTL catalogues offer the potential to gain insight into the functional consequences of the hundreds of coding and noncoding genetic loci that have been associated with neuropsychiatric disorders, including developmental delay (DD), ASD, SCZ, major depressive disorder (MDD), and Alzheimer’s disease (AD) (Deciphering Developmental Disorders Study, 2017; Grove et al., 2017; Kosmicki et al., 2016; Sanders et al., 2015; Sanders et al., 2017; Satterstrom et al., 2018; Schizophrenia Working Group of the Psychiatric Genomics Consortium, 2014). Many of these variants are presumed to act by altering gene expression, including noncoding SNPs that alter expression regulation, copy number variants (CNVs) that delete one copy of a gene, and premature stop codon variants that are predicted to induce nonsense-mediated decay (NMD) of transcripts (Nagy and Maquat, 1998; Rivas et al., 2015).

To help fill this gap, we have generated BrainVar, a unique resource of whole-genome sequencing (WGS) paired with bulk tissue RNA sequencing (RNA-seq) analyses of 176 samples from the human dorsolateral prefrontal cortex (DLPFC) across the span of its development, starting from 6 postconceptional weeks (PCW) to young adulthood (20 years). We focused our analyses on DLPFC due to its importance in higher order cognition (Silbereis et al., 2016) and the co-expression of many ASD and SCZ risk genes observed in this region during mid-fetal development (Gulsuner et al., 2013; Li et al., 2018; Network and Pathway Analysis Subgroup of the Psychiatric Genomics Consortium, 2015; Parikshak et al., 2013; Willsey et al., 2013). The single-base resolution captured by WGS across almost the entire genome allowed us to go beyond common SNPs captured by genotyping arrays used in previous eQTL studies of the human brain and detect other important classes of genetic variants, including those which span multiple basepairs (insertion/deletions (indels), copy number variants (CNVs)) or those that are rare or even unique to a person’s genome. Combined with high-depth paired-end RNA-seq in the same tissue samples, this approach allows hitherto unattainable analyses of the developmental transcriptome and the impact of diverse classes of genetic variants identified by WGS.

Our analysis discovers gene-specific developmental trajectories, identifying the genes that underlie the late-fetal transition, a dramatic shift in gene expression between mid-fetal development and infancy (Li et al., 2018). We characterize distinct groupings of genes as defined by their developmental expression trajectories and their co-expression patterns, and then leverage WGS to identify common variants implicated in cis-regulation of gene expression across development or during specific stages, and rare protein-truncating variants with corresponding changes in allele-specific and absolute gene expression. Finally, we evaluate relationships between these variants and genes associated with genome-wide significant loci in ten central nervous system (CNS) traits and disorders to identify unique temporal, cell type, and eQTL signatures.

## Results

### Description of the cohort and data generation

To characterize gene expression across prenatal and postnatal development of human DLPFC and identify genetic variants associated with expression changes, *post mortem* tissue was obtained from 176 de-identified, clinically unremarkable donors (genotypic sex: 104 male, 72 female) without known neuropsychiatric disorders or large-scale genomic abnormalities, ranging between 6 PCW and 20 years of age (Figure 1 and Table S1). In keeping with prior analyses (Kang et al., 2011), we assign these samples to 12 developmental stages, which we group into four developmental epochs (Figures 1 and 2). Gene expression data were generated using RNA-seq from tissue dissected from the frontal cerebral wall from donors aged <10 PCW or the DLPFC (corresponding mainly to Brodmann area 46) from donors at later developmental stages. WGS data (31.5x median coverage) were generated simultaneously from DNA isolated from the same individuals.

**Figure 1:**
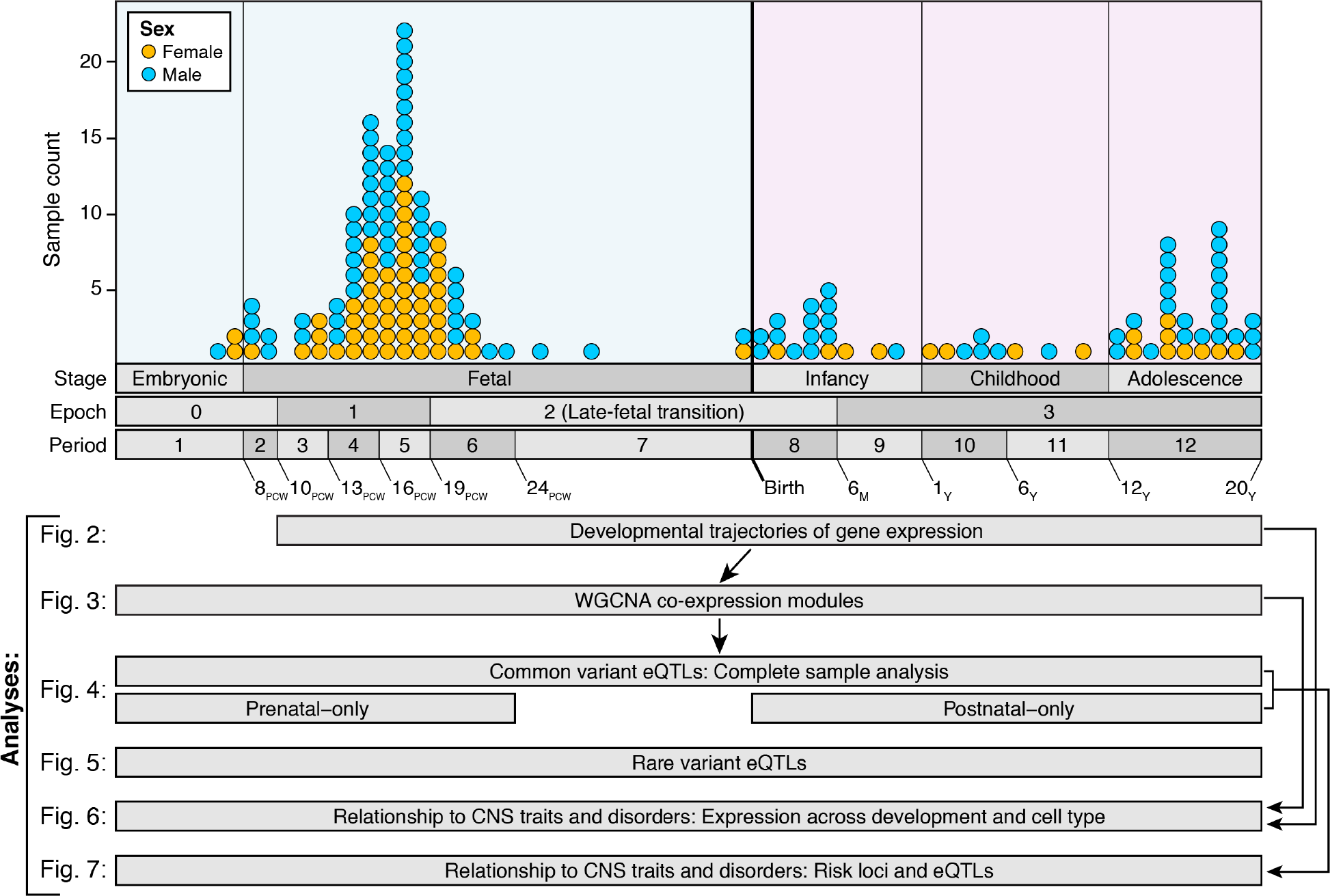
Overview of the dataset and the analysis. 176 samples from the dorsolateral prefrontal cortex (DLFPC) of the developing human brain were processed to generate RNA-seq gene expression data and WGS data (top). The distribution of the samples is shown by sex (color) and developmental stage (x-axis). Periods were defined previously (Kang et al., 2011) and epochs are defined as a superset of periods, based on principal component analysis of these RNA-seq data. The analyses conducted using these data are shown below, and the width of each box corresponds to the age of the samples included in each analysis.

**Figure 2.**
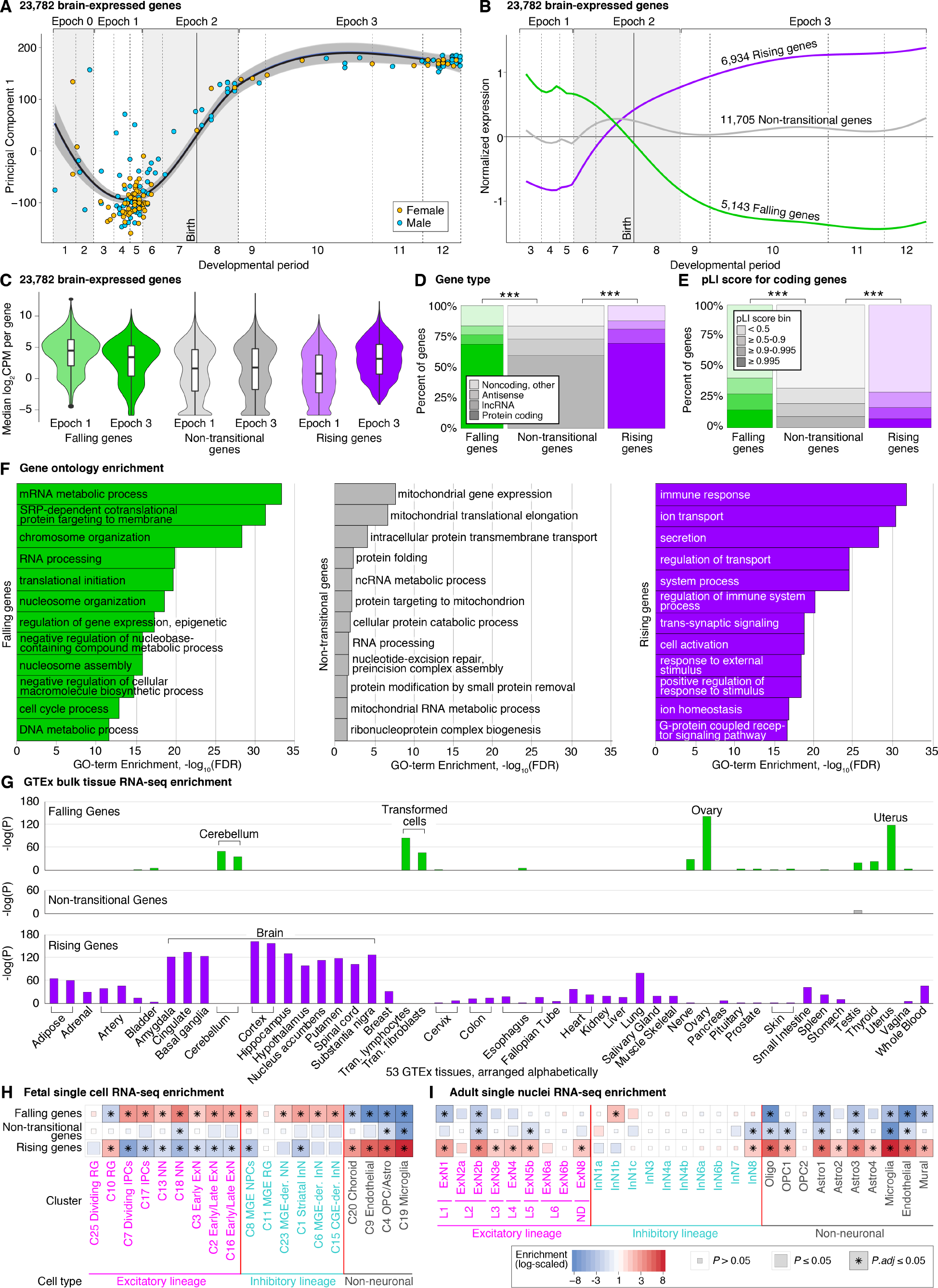
Temporal trajectories of gene expression in the human dorsolateral prefrontal cortex. **A)** Gene expression log_2_CPM for each sample was used to calculate principal components (Figure S2). The first principal component (PC1) explains 42% of the variance between samples and 81% of variance in PC1 is explained by developmental stage (Figure S2). The changes in PC1 over time were used to define four “epochs” of gene expression. Dotted lines represent the boundaries of the indicated developmental period as defined previously (Kang et al., 2011) **B)** Trajectory analysis was used to identify sets of genes with similar developmental profiles. Three groups of genes were identified and distinguished by their behavior across the late-fetal transition in epoch 2 (Table S2). For each group, the expression over time, normalized by the interquartile range and smoothed using locally weighted smoothing (LOESS) is shown. These three groups are further characterized by plotting: **C)** The median log_2_CPM across all samples in epoch 1 (E1) and epoch 3 (E3). **D)** The relative proportion of Gencode coding and noncoding genes. **E)** The distribution of probability loss-of-function intolerance (pLI) scores for protein-coding genes (Lek et al., 2016). **F)** The twelve gene ontology biological processes with the lowest FDR values. **G)** Enrichment in the most tissue-specific genes from the 53 tissues with bulk tissue RNA-seq data from GTEx. **H)** Enrichment of the 200 most cell type cluster-specific genes from 19 clusters defined by scRNA-seq data in the developing human cortex (Nowakowski et al., 2017). **I)** Enrichment of the 200 most cell type cluster-specific genes in 29 clusters defined by snRNA-seq data in the adult human DLPFC (Li et al., 2018). Abbreviations: CPM: Counts per million; RG: radial glia; IPC: intermediate progenitor cell; NN: newborn neuron; ExN: excitatory neuron; InN: inhibitory neuron; CGE: caudal ganglionic eminence; MGE: medial ganglionic eminence; OPC: oligodendrocyte precursor cell. Statistical analyses: A: Principal component analysis; B: Longitudinal mixture model with Gaussian noise; C: Wilcoxon two-sided; D: FET; E: gProfiler (Reimand et al., 2011); F: t-test with Benjamini-Hochberg correction; G: FET with Benjamini-Hochberg correction; H: FET, corrected for 57 comparisons; I: FET, corrected for 87 comparisons.

### Data processing

RNA-seq reads were aligned to GRCh38 using the STAR aligner (Dobin et al., 2013) with genes defined by Gencode v21 (Harrow et al., 2012). Read counts per gene were calculated using HTSeq (Anders et al., 2015) and converted to counts per million (CPM), which were logarithmically scaled to base 2 (log_2_CPM). Of the 60,155 genes assessed, 23,782 were defined as being cortically-expressed, based on CPM ≥ 1 in at least 50% of samples of either sex in at least one of the 12 developmental periods. We restricted further analysis to these 23,782 genes, for which median log_2_CPM ranges from −5.9 to 12.3, with a median of 2.3. Of these, 16,296 (68.5%) encode proteins, while 7,486 (31.5%) are noncoding, including lncRNA (12.6%) and antisense (9.2%) genes (Table S2). On average, noncoding genes were expressed at lower levels than protein-coding genes (3.1 log_2_CPM vs. −0.9 log_2_CPM; p<2×10^−16^, Wilcoxon test), especially lncRNAs (−1.5 log_2_CPM). However, not all noncoding genes follow this pattern, with 1,081 noncoding genes expressed above the median for all cortically-expressed genes and the nuclear lncRNA *MALAT1* being the fifth most highly expressed gene.

Although the vast majority of samples were newly ascertained for this study (162/176, 92%), 14 samples were previously analyzed by the BrainSpan Consortium (Li et al., 2018), but resequenced in BrainVar. For these 14 samples with replicate RNA-seq data in BrainSpan DLPFC samples, we compared the correlation of gene expression values with the current data. Despite differences in library preparation, with BrainSpan using poly-A priming compared to TruSeq random priming in BrainVar, the per sample correlations range from 0.76 to 0.91, while the median gene to gene correlation was 0.54 (Figure S1). The first principal component of gene expression in both data sets correlated strongly with developmental age (Figure S1).

WGS data were processed using a pipeline functionally equivalent to major WGS datasets (Regier et al., 2018), using BWA (Li and Durbin, 2009) for alignment to GRCh38, Picard (https://github.com/broadinstitute/picard/) to flag duplicate reads, and GATK v3.8 Haplotype Caller (McKenna et al., 2010) to call variants. Ancestry was estimated based on principal component analysis of 118,849 common SNPs and indels (≥5% minor allele frequency), alongside corresponding WGS data for 3,804 individuals of known ancestry (An et al., 2018). The results correlated strongly with self-reported ancestry and identified clusters corresponding to African-American ancestry (42 samples, 24% of cohort), European ancestry (82 sample, 47%), and Asian ancestry (4 samples, 2%), while 48 samples (27%) were outside of these clusters and enriched for individuals who identified as Hispanic, Alaskan native, or mixed ancestry (Table S1). Ancestry groups were differentially represented across developmental periods (F=3.4, df=5, p=0.006, ANOVA), with posthoc analysis showing a greater representation of African American samples in later developmental periods compared with samples of Hispanic, Alaskan native, or mixed ancestry (median age across cohort: one postnatal month vs. 17 PCW respectively; p.adj = 0.02, TukeyHSD).

To confirm concordant sample identity between the WGS and RNA-seq samples, we determined genetic sex, based on expression of *XIST* and chromosome Y genes, and sample genotype for 289 high fixation index (F_ST_) protein-coding SNPs (Table S1). All 176 samples had concordant sex and genotypes between the expression and sequencing data.

### Temporal dynamics of gene expression

Across the 176 samples, the first principal component explains 42% of the variance in gene expression and is highly correlated with developmental age (partial R^2^ for Period with PC1 = 0.88; Figure 2A and S2); the same pattern is observed for the 162 brain samples that are new to this analysis. This result is in keeping with prior analyses, highlighting the dramatic shift in gene expression across developmental age (Kang et al., 2011; Li et al., 2018). Similar to earlier reports, we also identified a coordinated shift in gene expression across multiple genes between mid-fetal development and early infancy (periods 6 to 8; epoch 2; Figure 2A), which we will refer to as the “late-fetal transition.” The data suggest a similar coordinated shift in embryonic and early fetal development (periods 1-2; epoch 0; Figure 2A). However, due to the sparse sampling (nine individuals), we did not characterize this “early-fetal transition” in detail. Using trajectory analysis, and excluding the nine epoch 0 individuals, we identified three distinct gene expression trajectories based on the presence or absence of persistent, progressive, and statistically significant expression variance across this late-fetal transition (Figure 2B). The groups are composed of 6,934 “Rising” genes, with expression rising through late-fetal development and infancy leading to higher postnatal expression; 5,143 “Falling” genes, with higher prenatal expression that then falls across late-fetal development leading to lower postnatal expression; and 11,705 “Non-transitional” genes that exhibit no statistically significant change in gene expression between mid-fetal development and childhood, adolescence, and/or adulthood. Emphasizing the magnitude of this transition, the first principal component of 11,705 genes with a Non-transitional trajectory explains only 18.4% of the variance in gene expression and is weakly correlated with developmental age (partial R^2^ for Period with PC1 = 0.3; Figure S2).

### Characteristics of transitional and non-transitional genes

To further characterize the nature of the late-fetal transition, we assessed the characteristics of the genes within the Rising, Falling, and Non-transitional groups. On average, the Falling genes were expressed at a higher level than both the Rising and Non-transitional groups throughout development, even in epoch 3, when their expression was lowest (3.4 vs. 3.2 median log_2_CPM for Falling vs. Rising genes in epoch 3; p=0.003, Wilcoxon test; Figure 2C). Falling genes had a higher proportion of coding genes than the Non-transitional genes (73.9 vs. 63.3%, p<1×10^−40^, chi-squared; Figure 2D) and were also enriched for genes with high probability Loss-of-function Intolerant scores (pLI ≥ 0.995; 10.1% of Falling genes, 1.6-fold expectation; Figure 2E), which identify genes with fewer protein truncating variants than expected in exome sequencing of ~60,000 population samples (Lek et al., 2016), suggesting haploinsufficiency resulting in selective pressure.

Gene ontology analysis of the Falling genes demonstrated enrichment for biological processes relating to development and anabolism, including cell division, gene expression regulation, RNA synthesis and processing, and protein translation (Figure 2F). Comparison with genes enriched in specific adult tissues from the Genotype-Tissue Expression (GTEx) project dataset (The GTEx Consortium, 2015) showed enrichment for several non-cortical tissues (Figure 2G), driven by genes with ontology terms relating to RNA transcription in the cerebellum, ovaries, and uterus and cell division in transformed cells (Table S2). Compared to H3K27ac DLPFC peaks from BrainSpan (Li et al., 2018), Falling genes were enriched in genes associated with fetal-biased H3K27ac peaks (Figure S2). To further understand how these temporal changes relate to cell differentiation and cell type, we assessed the overlap of our bulk tissue expression with data from single-cell RNA-seq (scRNA-seq) in prenatal human forebrain (Nowakowski et al., 2017) and single-nucleus RNA-seq (snRNA-seq) in postnatal DLPFC (Li et al., 2018). Using cell clusters previously defined in each of these studies, we applied a Tau metric to define the 200 protein-coding genes most enriched in each cell type cluster. The Falling genes are enriched in multiple neuronal cell type clusters of the excitatory and inhibitory lineages, mainly from the prenatal data (Figure 2H).

The Rising genes had median postnatal expression values that were lower than Falling genes (3.2 vs. 3.4 median log_2_CPM in epoch 3), though a similar proportion of protein-coding genes (73.1 vs. 73.9%). The Rising genes were enriched for adult-biased H3K27ac peaks (Figure S2) but depleted for genes with high pLI scores (5.1% pLI ≥ 0.995; 0.80-fold expectation), suggesting greater tolerance to variation in expression. Gene ontology terms enriched in the Rising genes related to immune processes and metabolic processes, such as transportation, secretion, and ion homeostasis, with the genes highly enriched in all adult brain tissues in GTEx except the cerebellum. In the single cell data, the Rising genes are enriched in multiple non-neuronal cell types, including microglia, oligodendrocyte precursor cells (OPC), oligodendrocytes, and astrocytes in both the pre- and postnatal data, and multiple classes of excitatory neurons, mainly in the postnatal data (Figure 2I).

The Non-transitional genes had the lowest levels of expression, the lowest proportion of protein-coding genes, and few genes with high pLI score (5.4%, 0.86-fold expectation). They were not enriched for either fetal-biased or adult-biased H3K27ac peaks (Figure S2), mapped to gene ontology terms relating to mitochondria and protein folding and transport and were expressed ubiquitously across GTEx tissues. None of the cell type clusters in pre-or postnatal scRNA-seq data were enriched for Non-transitional genes and, compared to Transitional genes (Falling and Rising), the Non-transitional genes were less likely to be enriched in a cell type cluster (14.3% of Transitional genes vs. 8.3% of Non-transitional genes; odds ratio (OR) = 1.8 (95%CI: 1.7-2.0); p≤2×10^−16^ FET).

### Co-expression modules in the developing human cortex

To further define the relationships between the 23,782 cortically-expressed genes, we applied weighted gene co-expression network analysis (WGCNA) to define 19 consensus modules that included 10,459 genes (Figures 3A and S3). As expected, genes within each module shared functional roles (Figure 3B), temporal trajectories of gene expression (Figure 3C and 3D), and regulatory transcription factors (Figure S3). In contrast, genes that were not assigned to modules were more likely to be noncoding (OR = 2.5; 95%CI: 2.3-2.6; p<2×10^−16^, FET), have lower expression (0.74 vs. 3.20 median log_2_CPM; p<2×10^−16^, Wilcoxon test), and belong to the Non-transitional group (OR = 1.7; 95%CI: 1.6-1.8; p<2×10^−16^, FET). Module preservation analysis using BrainSpan data (Li et al., 2018) identifies similar co-expression patterns in independent DLPFC samples and across all neocortical samples, but not in non-neocortical regions (Figure 3E). Comparison of the modules to the prenatal and postnatal scRNA-seq data showed enrichment for specific cell type clusters for ten of the modules, with patterns similar to those observed for the temporal trajectories (Figure 3F and 3G).

**Figure 3.**
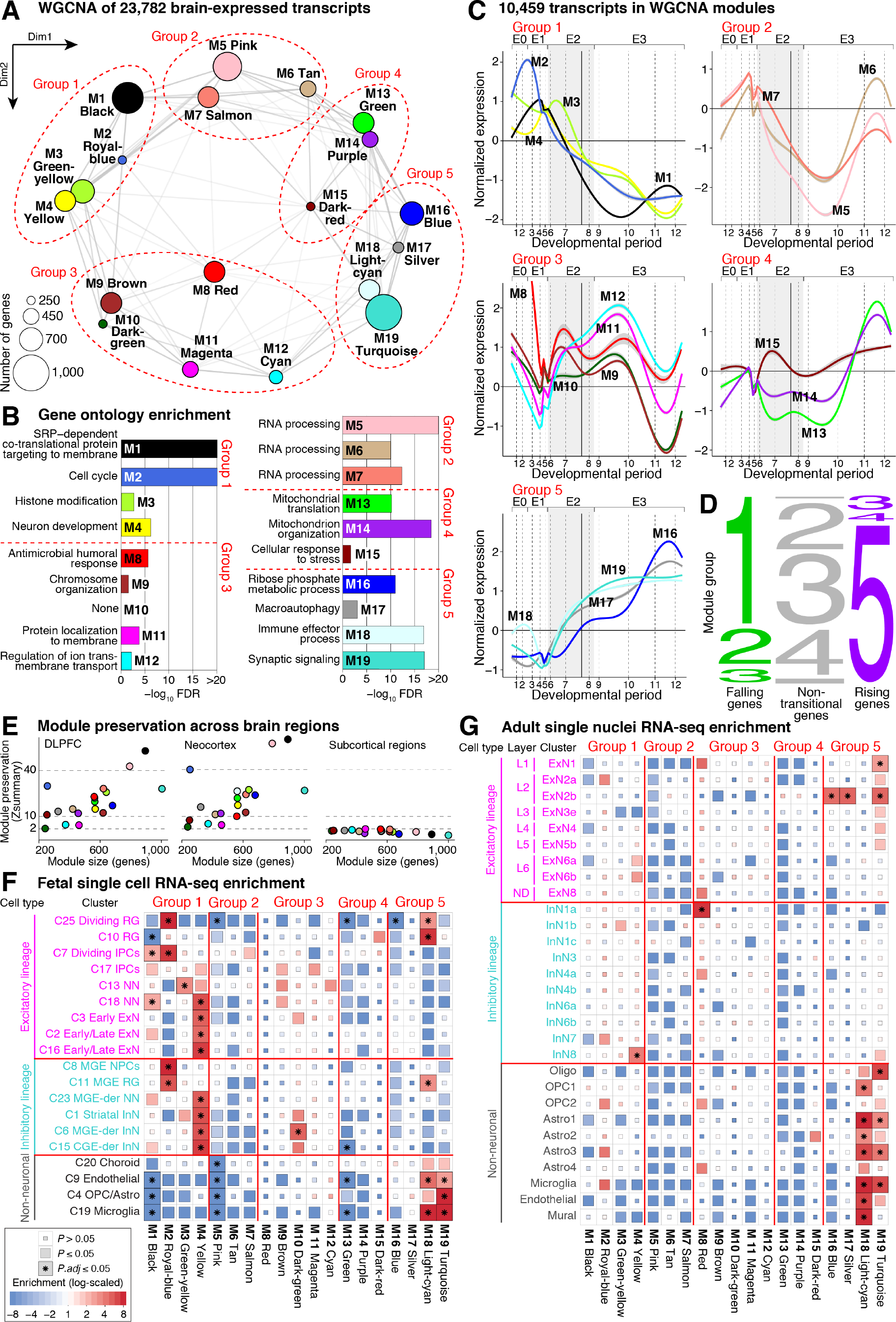
Co-expression modules in the developing human cortex. **A**) WGCNA identified 19 modules comprised of 10,459 genes, shown as colored nodes, from patterns of co-expression for 23,782 expressed genes. The nodes are plotted based on the first two dimensions from multidimensional scaling. The weight of the connecting lines represents the degree of correlation between module eigengenes. **B)** Normalized expression is shown for the 19 modules arranged in five groups based on proximity in ‘A’ and similar temporal trajectories. **C)** Gene ontology enrichment analysis for each module, showing only biological processes with the lowest FDR; M10 Dark-green, was not enriched for any process after correcting for multiple comparisons. **D)** The relationship between the five groups of co-expression modules (from ‘A’) and the genes with Falling, Rising, or Non-transitional temporal trajectories (Figure 2). The number represents the WGCNA group, with the height proportional to the number of genes in the group and the width proportional to the number of genes in each temporal trajectory group. Detailed relationships between modules and temporal trajectories are shown in Figure S3. **E)** Module preservation in independent BrainSpan samples (Li et al., 2018) from the same brain region (left), other cortical regions (middle), and five subcortical regions (right). **F)** Enrichment between the 19 modules and the 200 genes most specific to 19 cell type clusters defined by scRNA-seq data in the developing human cortex (Nowakowski et al., 2017). **G)** Enrichment between the 19 modules and the 200 genes most specific to 29 cell type clusters defined by snRNA-seq data in the adult human DLPFC (Li et al., 2018). Abbreviations: FDR: False discovery rate; SRP: Signal Recognition Particle. Statistical analysis: A: WCGNA with the consensus module detection from 100 random resampling; B: FET, corrected for gProfiler gene ontology pathways (10 to 2000 term size); C: gene expression normalized by interquartile range; E: FET, corrected for 361 comparisons; F: FET, corrected for 551 comparisons. G: FET, corrected for 19 comparisons.

In keeping with our trajectory analysis (Figure 2), multidimensional scaling analysis of the module eigengenes demonstrated that developmental age accounted for 44.7% and 36.1% of the variance in the first two dimensions. Plotting the relationship of 19 modules between these two dimensions, and considering the developmental trajectories of the genes in each module, we identified five groups of related modules (Figure 3A and 3B). Group 1 modules (M1 Black, M2 Royal-blue, M3 Green-yellow, M4 Yellow) are closely related to the Falling genes (Figure 3D) and have similar characteristics, including enrichment for gene ontology terms related to cell division, protein translation and targeting, and histone modifications, along with enrichment in developing neurons of the excitatory and inhibitory lineages. Of note, the M4 Yellow module is enriched for numerous ontology terms related to neuronal development, contains genes specific to neuronal stem cells (e.g. *NCAM1*/*ncam*, *PROM1*/*cd133*), and is highly enriched for genes related to maturing excitatory and inhibitory neurons, while the M2 Royal-blue module, which captures cell cycle, is enriched in neuroprogenitor cells, radial glia, and intermediate progenitor cells. Group 2 modules (M5 Pink, M6 Tan, M7 Salmon) are enriched for factors relating to RNA processing (Figure 3C) and contain targets of numerous transcription factors and microRNAs (Figure S3). Gene expression of group 2 modules increases between periods 10-11 (1-12 years post birth), suggesting a key role for RNA processing during childhood cortical development.

Group 3 modules (M8 Red, M9 Brown, M10 Dark-green, M11 Magenta, M12 Cyan) overlap with the Non-transitional gene set, with expression peaking between periods 6-9 (19 PCW to 1-year post birth) before returning to mid-fetal levels. They are enriched for terms related to protein localization, ion transport, synapses, and receptors. The M8 Red module, which is enriched for ontology terms relating to cell fate and morphogenesis, is highly enriched for noncoding genes and has an expression peak in early fetal development, hinting at an early-fetal transition in epoch 0 analogous to the epoch 2 late-fetal transition. Several genes associated with regional patterning in non-cortical tissues, including hindbrain (e.g. *UNCX*, *CCDC140*) and hypothalamus (e.g. *DMBX1*, *SOX14*), are expressed at high levels in this module. Group 4 modules (M13 Green, M14 Purple, M15 Dark-red) are relatively static during development before rising steeply between periods 10 and 11 (1-12 years post birth), mirroring the group 2 modules. They are enriched for multiple ontology terms relating to mitochondria and cellular respiration, possibly reflecting the metabolic activity necessary to support neuronal activity during this postnatal stage (Ashrafi and Ryan, 2017). Finally, the group 5 modules (M16 Blue, M17 Silver, M18 Light-cyan, M19 Turquoise) are closely related to the Rising genes (Figure 2) with expression rising throughout development. Both the M18 Light-cyan and M19 Turquoise modules are strongly enriched in glial and other non-neuronal cell clusters, accordingly, both modules are enriched for ontology terms related to immune responses. The M19 Turquoise module is also enriched in excitatory neurons on the postnatal cortex and ontology terms relating to synaptic signaling and neurotransmitter transport. The other two modules in group 5, M16 Blue and M17 Silver, are enriched for terms relating to numerous metabolic processes, including glycolysis, respiration, and protein catabolism, along with neuronal death and autophagy.

### Common genetic variants regulating gene expression

We next leveraged WGS data to identify 6,573,196 high-quality SNPs and indels, observed at an allele frequency of at least 5% in both our prenatal (periods 1-6, N=112) and postnatal (periods 8-12, N=60) samples. The four samples from period 7 were excluded due to the sparse representation over this dynamic developmental phase (Figure 2). This common variant set included only variants located outside of low-complexity regions (Li, 2014) that passed variant quality score recalibration, with allelic balance between 0.22-0.78 (SNPs; 0.2-0.8 for indels), Hardy Weinberg equilibrium p-value ≥1×10^−12^, call rate ≥90% across all samples, and that reached variant and genotype thresholds established and validated in a previous analysis of family-based WGS data (Werling et al., 2018).

Linear regression was used to identify common variants within 1Mb of a gene associated with altered levels of gene expression, i.e. cis-eQTLs. Developmental period, sex, and the first five principal components for ancestry were included as covariates. Subsequently, we applied two approaches to control for technical covariates: Hidden Covariates with Prior (HCP) and surrogate variable analysis (SVA). To distinguish temporally specific eQTLs, we performed cis-eQTL analysis for three sets of samples: all 176 samples (complete sample), 112 samples in periods 1-6 (prenatal), and 60 samples in periods 8-12 (postnatal). Within each analysis, results for all tested variant-gene pairs were corrected for multiple testing using the Benjamini-Hochberg procedure.

Across these three analyses, we identified 252,629 unique eQTLs (FDR ≤ 0.05) associated with the expression of 8,421 unique genes (eGenes) (Table S4). While HCP-adjusted results were used for our primary analysis, results for 167,665 unique eQTLs identified using SVA are also provided (Table S4). The rate of eGene discovery from each analysis aligns with expectation given sample size (Figure S4).

We observed no significant difference in the effect size between SNVs and indels (p = 0.55, two-sided Wilcoxon test) or between insertions and deletions (p = 0.87, two-sided Wilcoxon test; Figure S4). Considering functional annotations, we identified greater overlap with H3K27ac peaks derived from the human brain (Li et al., 2018; Reilly et al., 2015) for eQTLs than non-eQTL variants (Figure S4). Analysis with GREGOR (Schmidt et al., 2015) using 18 human frontal lobe derived chromatin states (Roadmap Epigenomics Consortium et al., 2015) showed enrichment of eQTLs near transcription start sites and genic enhancers (Figure S4).

### Temporal specificity of eQTLs

Given the dramatic changes in gene expression across development, we leveraged the inclusion of both prenatal and postnatal samples to assess whether eQTLs also varied across development. To delineate the temporal specificity of eQTLs, we categorized eQTLs and their associated eGenes according to the magnitude and significance of their effects in our prenatal, postnatal, and complete analyses. We defined 161,293 (63.8% of 252,629 eQTLs) “Constant” eQTLs as those observed in all three analyses (FDR ≤ 0.05 in the complete sample, uncorrected p ≤ 0.05 in both the prenatal and postnatal samples) with the same direction of effect. The 19,142 (7.6%) “Prenatal-specific” eQTLs were observed in the prenatal data (FDR ≤ 0.05), but not the postnatal data (p > 0.05) and had a significant difference in effect size between these two cohorts (FDR-corrected Z-test of regression coefficients p ≤ 0.05). Analogous criteria were used to define 7,973 (3.2%) “Postnatal-specific” eQTLs. For the remaining eQTLs, the absolute value of the effect size in the prenatal and postnatal data sets was used to define 55,659 (22.0%) “Prenatal-trending” eQTLs and 7,931 (3.1%) “Postnatal-trending” eQTLs (Figure 4A).

**Figure 4:**
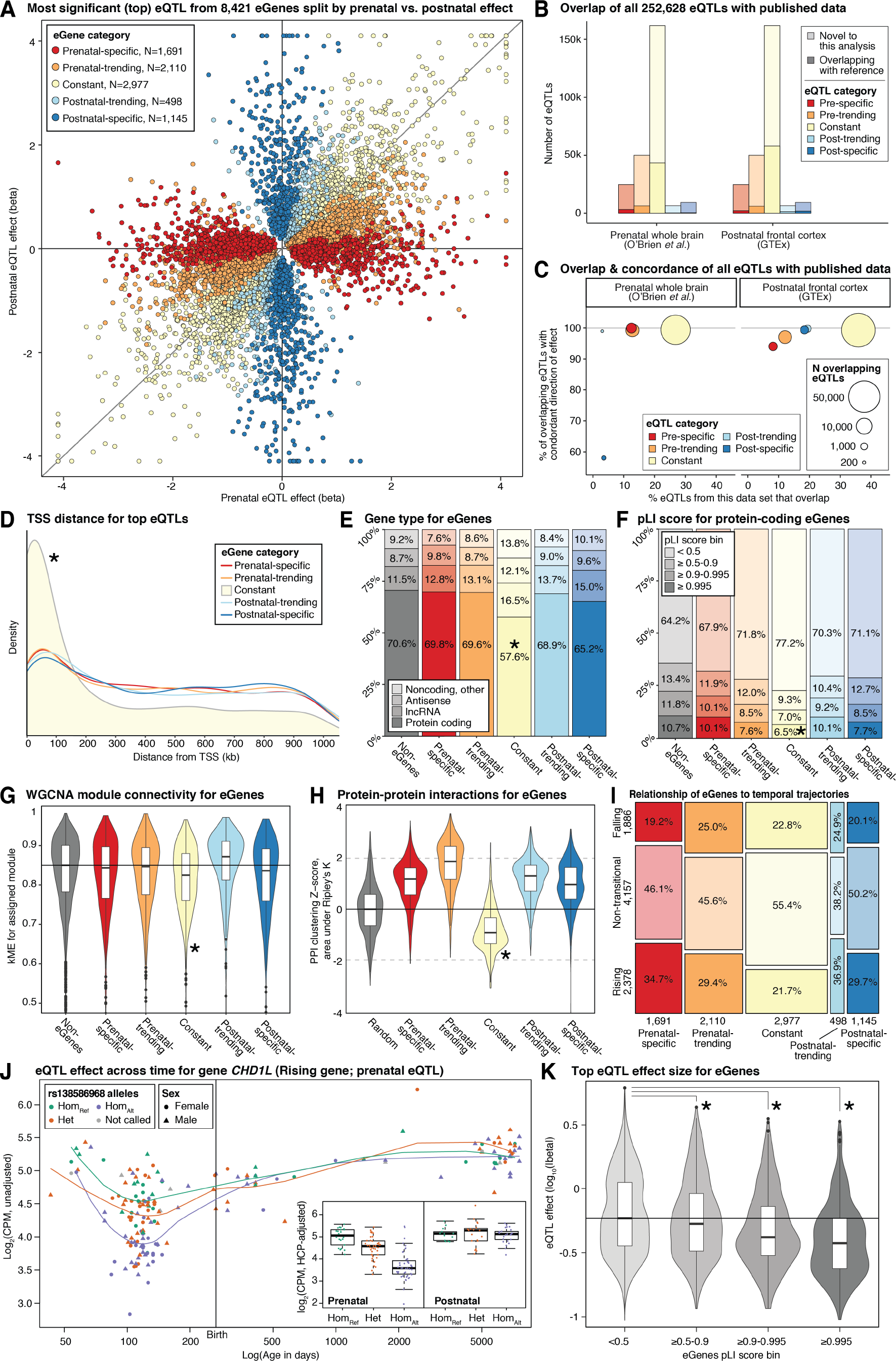
Common variant cis-eQTLs. **A)** Prenatal (x-axis) and postnatal (y-axis) effects for the eQTLs with the smallest p-value for 8,421 eGenes (points). The eQTLs are split into five categories based on temporal specificity using effect size and statistical thresholds; categories are represented by color. **B)** The total number of eQTLs in each temporal category is represented by the height of the bar. The number of eQTLs that match previously identified significant eQTLs in the prenatal whole brain (O’Brien et al., 2018) (left) and postnatal frontal cortex (The GTEx Consortium et al., 2017) (right) is shown by the darker shade. **C)** The percentage overlapping eQTLs between datasets (from ‘B’) is shown on the x-axis against the percentage of these overlapping eQTLs with concordant direction of effect in both datasets. **D)** Density plot of the distance of top eQTLs per eGene from the TSS by eGene temporal category. Characteristics of non-eGenes and eGenes in each temporal category are shown by plotting the: **E)** proportion of coding and noncoding genes, **F)** proportion of genes with pLI scores in different bins, **G)** centrality to an assigned WGCNA module (kME values, black line is the non-eGene median), and **H)** BioGRID protein-protein interactions (permuted Z-scores from Ripley’s K-net function (Cornish and Markowetz, 2014), black line is the non-eGene median). **I)** Mosaic plot of the proportion of genes in each temporal trajectory with eGenes split by temporal category. **J)** Expression data binned by genotype for the top Prenatal-specific eQTL for CHD1L, a gene with a Rising trajectory. Main panel: Gene expression by sample age across development. Lines are loess-smoothed trajectories for expression in samples with each of three genotypes for rs138586968. Inset: Boxplots for prenatal (left) and postnatal (right) samples with each of three rs138586968 genotypes. **K)** Distribution of eQTL effect size for eGenes binned by pLI scores, the black line is the median of the pLI < 0.5 group. Abbreviations: eQTL: expression quantitative trait locus; eGene: gene regulated by an eQTL; TSS: transcription start site; pLI: probability loss-of-function intolerant. Statistical analysis: D, G, H: two-sided Wilcoxon rank sum test for Constant versus other eGenes; E, F: two-sided FET for Constant versus other eGenes; K: two-sided Wilcoxon rank sum test for three top pLI bins versus genes with pLI < 0.5.

To assess concordance with comparable datasets, we assessed FDR-significant eQTLs that overlapped between our data and prenatal whole brain (O’Brien et al., 2018) or postnatal frontal cortex (The GTEx Consortium et al., 2017). The percentage of overlapping eQTLs was highest for the Constant group (26.8% with prenatal brain and 35.9% with postnatal frontal cortex; Figure 4B), while 73.2% and 64.1% (118,543 and 103,868 constant eQTLs), respectively, were novel to this analysis. Of the Constant eQTLs that did overlap, the direction of effect was concordant for 99.4% and 99.3% of eQTLs respectively, despite differing cohorts, library preparation methods, genome-build, the use of WGS, and analysis pipelines. As expected, temporally specific eQTLs showed higher overlap and concordance in samples of an equivalent developmental age (e.g. Prenatal-specific eQTLs in prenatal whole brain) compared to samples of a different developmental age (e.g. Postnatal-specific eQTLs in prenatal whole brain; Figure 4C). A comparison of the prenatal whole brain and postnatal cortex reference data sets to each other demonstrated comparable or a lower percentage of overlapping eQTLs (29.7% of prenatal brain eQTLs in postnatal frontal cortex; 13.9% of postnatal frontal cortex eQTLs in prenatal brain) than we attain for Constant eQTLs, with only 93.9% concordance in the direction of effect (Figure S4).

### Characteristics of genes influenced by eQTLs

More than one eQTL was identified for 5,538 of the 8,421 eGenes (65.8%), often with similar direction and magnitude of effect due to linkage disequilibrium. We therefore identified the top eQTL for each eGene, defined as the lowest FDR-significant p-value in any of the three sample sets (Table S4). By allowing the eGene to inherit the temporal classification of this top eQTL, we categorized 1,691 eGenes as Prenatal-specific, 2,110 as Prenatal-trending, 2,977 as Constant, 498 as Postnatal-trending, and 1,145 as Postnatal-specific (Figure 4A). The temporal-specific category of the top eQTL matched the majority of eQTLs in the gene for 7,425 (88.2%) of eGenes (Fig S4), with a further 688 (8.2%) of genes shifting by no more than one category (e.g. Prenatal-specific to Prenatal-trending, Postnatal-trending to Constant). While 63.8% of all eQTLs are Constant, only 3,804 (45.2%) eGenes have any Constant eQTLs. This difference is because many Constant eQTLs have high rates of linkage disequilibrium and regulate the same gene.

eQTLs are often observed in close proximity to the Transcription Start Site (TSS) of the associated gene (Fromer et al., 2016; The GTEx Consortium et al., 2017). This pattern was evident for the top eQTLs of the 2,977 Constant eGenes with a median distance of 92,223bp and both statistical significance and effect size of the eQTL increasing with proximity to the TSS (p = 6.6×10^−66^ for −log_10_(P), p = 1.0×10^−12^ for |*β*|, linear regression). In contrast, the median distance for the top eQTLs of the 5,444 eGenes with a degree of temporal specificity was 402,591bp (p = 1.3×10^−148^ two-sided Wilcoxon rank sum test compared to Constant eGenes, Figure 4D) with statistical significance, but not the effect size, of the eQTL increasing with proximity to the TSS (p = 2.8×10^−41^ for −log_10_(P), p = 0.25 for |*β*|, linear regressions for distance from TSS.

This distinction between Constant eGenes and those with a degree of temporal specificity was also observed across multiple other metrics. Compared to other eGenes, Constant eGenes include fewer protein-coding genes (OR = 0.62 (95%CI: 0.57-0.68), p = 6.5×10^−24^, two-sided Fisher’s exact test (FET); Figure 4E). Of the protein-coding genes, Constant eGenes also include fewer genes with high pLI scores than other eGenes (≥ 0.995; OR = 0.74 (95%CI: 0.58- 0.93), p = 0.01, two-sided FET; Figure 4F). Considering the WGCNA results, Constant eGenes are less likely to be assigned to a module (OR = 0.66 (95%CI: 0.61-0.73), p = 3.0×10^−18^, two-sided FET) and, for those that are assigned a module, show weaker co-expression with the module eigengene (0.83 vs. 0.85 median kME, p = 2.5×10^−9^, two-sided Wilcoxon rank sum test; Figure 4G). Constant eGenes also show weaker connectivity to PPI networks (median Z-score −0.914 for Constant vs. 1.294 for other eGenes, p = ≤2×10^−16^, two-sided Wilcoxon rank sum test; Figure 4H). Across these metrics, eGenes with a degree of temporal specificity tend to be more similar to non-eGenes than the Constant eGenes.

Temporally specific eQTLs could be a consequence of higher expression yielding better power to detect variation between samples at one developmental stage; this explanation would predict that Prenatal-specific eGenes would be enriched for Falling genes and Postnatal-specific eGenes would be enriched for Rising genes. We do not observe this effect (Figure 4I); instead, the Prenatal-specific eGenes are enriched for Rising genes (OR = 1.5 (95%CI: 1.3-1.6), p = 1.1×10^−10^, two-sided FET) and depleted for Falling genes (OR = 0.79 (95%CI: 0.69-0.9), p = 4.2×10^−4^, two-sided FET), while the distribution of Postnatal-specific eGenes across trajectories does not differ significantly from expectation. The Rising gene *CHD1L* and Prenatal-specific eQTL rs138586968 provides a representative example of this pattern (Figure 4J).

Comparing eGenes to WGCNA modules (Figure S4), we find that Constant eGenes are enriched for four of the five Group 3 modules, with significant enrichment for the M9 Brown module (OR = 2.1 (95%CI: 1.6-2.8), p = 1.46×10^−6^, two-sided FET); Constant eGenes are also significantly depleted from the M19 Turquoise module (OR = 0.38 (95%CI: 0.27-0.53), p = 1.1×10^−8^, two-sided FET) and the M8 Red module (OR = 0.47 (95%CI: 0.32-0.69), p = 2.83×10^−3^, two-sided FET). Postnatal-specific eGenes are enriched for the M8 Red module (OR = 2.5 (95%CI: 1.8-3.5), p = 1.77×10^−5^, two-sided FET; Figure S4), which is characterized by high expression in epoch 0 and low expression thereafter (Figure 3; Table S2), and Prenatal-specific eGenes are depleted for the M3 Green-yellow module (OR = 0.51 (95%CI: 0.33-0.76), p = 0.04, two-sided FET), which has high prenatal and lower postnatal expression.

The lack of enrichment for Prenatal-specific eGenes in the Group 1 modules and the Falling genes or Postnatal-specific eGenes in the Group 5 modules and the Rising genes raises the possibility that natural selection constrains the variability of gene expression at developmental stages where the genes perform critical functions, necessitating high expression. We reasoned that selective pressure would similarly constrain the variability in gene expression of eQTLs near dosage sensitive genes. Accordingly, we observed an inverse relationship between eQTL effect size and pLI score (0.59 vs. 0.38 median |*β*| for genes with pLI < 0.5 vs. pLI ≥ 0.995-1; p = 4.0×10^−36^; two-sided Wilcoxon rank sum test; Figure 4K).

### Allele specific expression and eQTLs for rare variants

Along with identifying common variant eQTLs, WGS data raises the possibility of detecting rare variants that alter gene expression. However, at allele frequencies below 5%, rare variants would be present in eight or fewer samples in our cohort of 176, limiting the power for detection in a genome-wide analysis. We therefore focused on rare protein truncating variants (PTVs) with a premature stop codon and exonic deletions in cortically expressed protein-coding genes predicted to halve gene expression. To identify the rarest variants, which have the potential to mediate the highest effects, we defined rare for PTVs as never observed in 15,708 individuals with WGS in the Genome Aggregation Database (GnomAD; (Lek et al., 2016) or in 3,804 unrelated samples with WGS data from the Simons Simplex Collection (SSC) and, for deletions, never observed in 9,097 samples from SSC families (An et al., 2018; Werling et al., 2018).

We expect a PTV to induce nonsense-mediate decay (NMD) of the mRNA, resulting in allele specific expression, i.e. the absence of reads containing the variant in RNA-seq data (Nagy and Maquat, 1998; Rivas et al., 2015). We identified 88 rare PTVs in Gencode-defined protein-coding genes with at least five RNA-seq reads overlaying the variant and classified them into three categories: 22 “Escaping NMD” variants in the last coding exon or 50bp upstream of the last splice site (Nagy and Maquat, 1998); 11 “Possible NMD” variants in exons that are not expressed in all isoforms or in single-exon genes; and “Probable NMD” for the remaining 55. For the Probable NMD group, 45 of the 55 PTVs (81.8%) had fewer than 50% of reads that contained the variant (p=1×10^−6^, binomial exact test; Figure 5A), with a mean of 35.3% (95% CI: 30.4 to 40.2%), which was lower than the mean of 48.8% (95% CI: 44.2 to 53.5%; p=0.001, Wilcoxon test) in the Escaping NMD group, in which 11 of 22 PTVs (50.0%) had fewer than 50% variant reads (p=0.58, binomial exact test). The Possible NMD group is similar to the Probable NMD group (10 out of 11 had below 50% variant reads, p=0.006, binomial exact test; mean of 31.3%, 95% CI: 27.9 to 34.6%).

**Figure 5.**
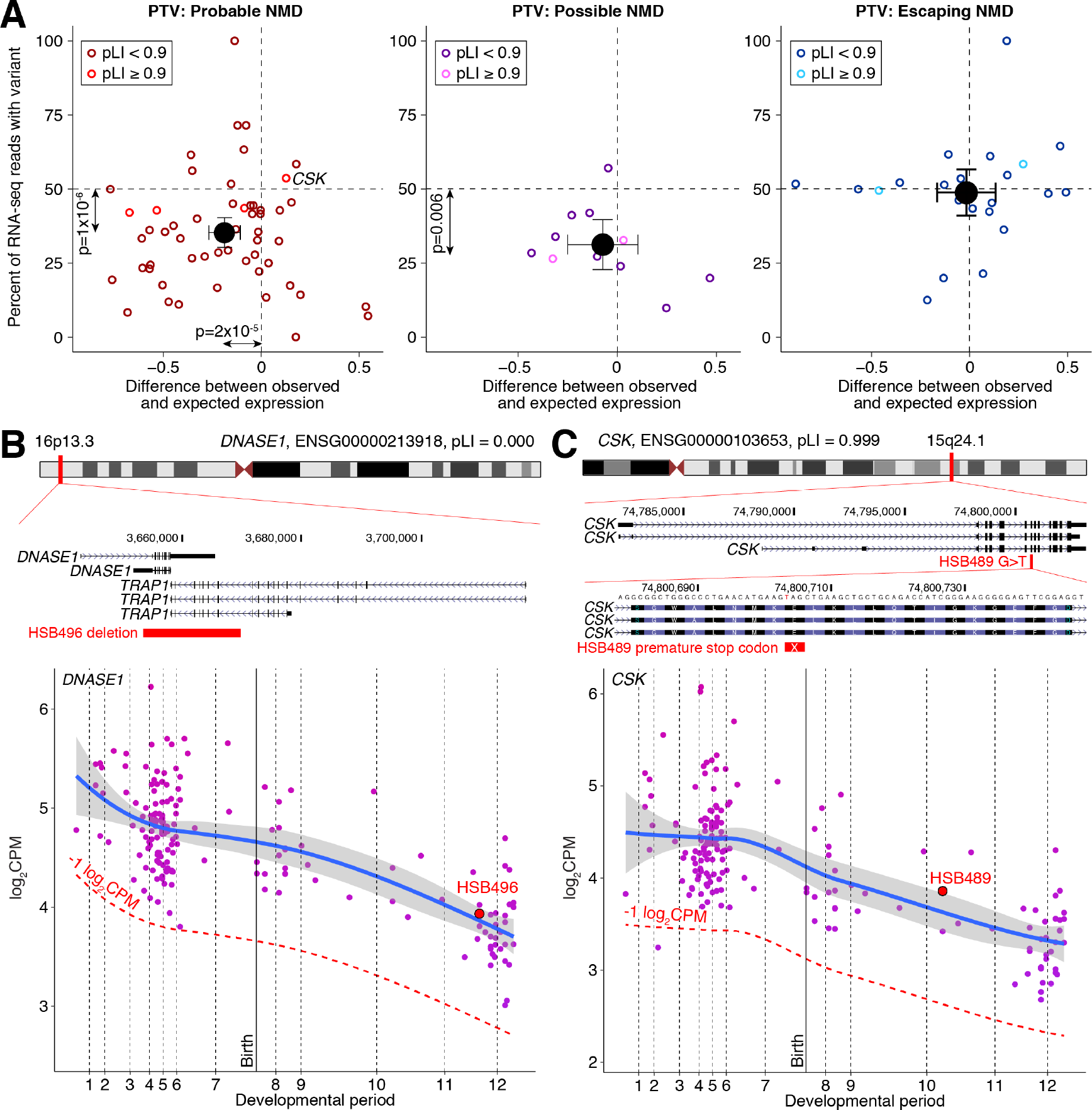
Rare variant eQTLs. **A)** For 55 rare PTVs (red points) predicted to undergo NMD (Probable NMD) with at least 5 RNA-seq reads at the site of the variant, the percentage of RNA-seq reads containing the variant (y-axis) is plotted against the difference between the observed and imputed log_2_CPM in the individual with the PTV (x-axis). The black point shows the centroid and 95% confidence intervals across all 55 PTVs. This plot is repeated for 12 “Possible NMD” PTVs in single exon genes or with the PTV-containing exon varying between isoforms (middle) and 22 “Escaping NMD” PTVs in the last exon or 50bp upstream of the last splice site (right) (Nagy and Maquat, 1998; Rivas et al., 2015). **B)** A deletion of all protein-coding exons in the Deoxyribonuclease 1 (DNASE1) gene and the first seven protein-coding exons of TNF receptor associated protein 1 (TRAP1) in sample HSB496 has no appreciable effect on the expression level of DNASE1 (shown) or TRAP1 (Figure S5). **C)** A variant causing a premature stop codon in a constitutive exon in the C-Terminal Src Kinase (CSK) gene in sample HSB489 has no appreciable effect on expression level or allele specific expression (53.7% variant reads), despite being predicted to cause NMD. Abbreviations: PTV: Protein truncating variant; NMD: Nonsense-mediated decay; pLI: probability loss-of-function intolerant. Statistical analysis: A: binomial exact test.

Alongside allele specific expression, we also expect NMD to induce a 50% reduction in gene expression, equivalent to −1 log_2_CPM. To evaluate this expectation, we assessed the difference between observed and expected expression, with the later estimated by imputation accounting for variation between samples and across developmental periods. In the Probable NMD group, expression was lower than expectation for 43 of the 55 PTVs (p=2×10^−5^, binomial exact test), with a mean of −0.19 log_2_CPM (95%CI: −0.11 to −0.27), equivalent to a 12.1% (95%CI: 7.2 to 16.9%) reduction in expression. This was also lower than the mean of −0.02 log_2_CPM observed in the Escaping NMD group (p=0.02, Wilcoxon test). For genes with lower median log_2_CPMs, and consequently fewer reads from which to estimate expression, we observed greater variability in gene expression (Figure S5). Of note, none of the Probable NMD PTVs resulted in the expected magnitude of decreased expression, with the largest difference being −0.77 log_2_CPM, equivalent to a 41% reduction. In contrast, we observed numerous examples of variants predicted to decrease expression that did not, including a heterozygous deletion of all protein-coding exons in the *DNASE1* gene (pLI = 0.000; Figure 5B, S5) and a premature stop codon in the haploinsufficient *CSK* gene (pLI = 0.999; Figure 5C).

### Temporal trajectories in human traits and disorders

To leverage this dataset to provide insight into the biology underlying CNS traits and disorders, we assessed the enrichment of gene lists based on developmental trajectory, module, and cell type for genes associated with human traits and disorders. We selected ten datasets that applied genome-wide statistical thresholds to identify multiple associated variants. For DD, ASD, and epilepsy with developmental delay (EPDD) we used gene lists derived from rare and *de novo* variants identified with exome sequencing (Deciphering Developmental Disorders Study, 2017; Heyne et al., 2018; Satterstrom et al., 2018). For educational attainment (EA), attention deficit/hyperactivity disorder (ADHD), SCZ, MDD, multiple sclerosis (MS), Parkinson’s Disease (PD), and AD we used genes within 10kb of common variants detected with genotyping arrays in genome-wide association studies (GWAS) (Chang et al., 2017; Demontis et al., 2019; International Multiple Sclerosis Genetics Consortium et al., 2013; Lambert et al., 2013; Lee et al., 2018; Schizophrenia Working Group of the Psychiatric Genomics Consortium, 2014; Wray et al., 2018). For our analyses, we excluded genes within the Major Histocompatibility Complex on chromosome 6, due to the complicated nature of this region (Schizophrenia Working Group of the Psychiatric Genomics Consortium, 2014).

To assess temporal relationships for CNS traits and disorders we plotted the normalized gene expression across development, ordering the disorders by the mean period of maximal expression for each gene (Figure 6A), and assessed the enrichment for the transitional gene groups (Figure 6B). In general, early-onset traits and disorders, including DD, EA, and ASD, have higher prenatal than postnatal expression and are enriched for Falling genes. In contrast, later-onset disorders, including PD and AD, have higher postnatal expression and are enriched for Rising genes. EPDD would be expected to group with DD if there were a simple relationship between maximal gene expression and the onset of a disorder, however it is ranked last on this list, after PD and AD. We note that, while GWAS loci are genome-wide significant, the genes associated with these loci are likely to contain false positives, which may obscure clearer temporal relationships. Furthermore, critical periods of neurological vulnerability may precede symptom onset.

**Figure 6.**
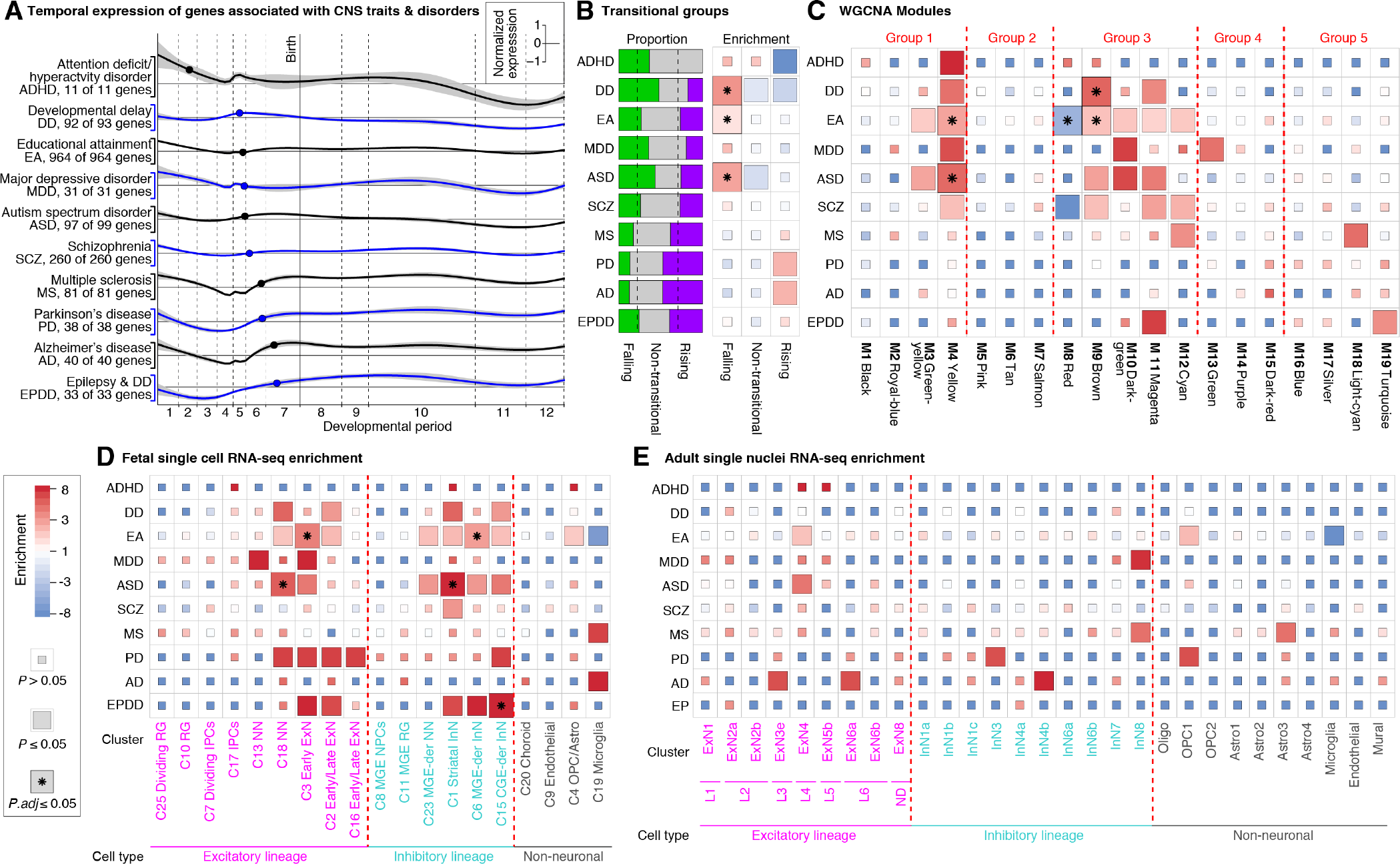
Expression of genes associated with CNS traits and disorders. **A)** Genome-wide significant loci were collated for ten CNS traits and disorders from exome sequencing (DD, ASD, EPDD) or GWAS (ADHD, MDD, EA, SCZ, MS, PD, AD) and genes associated with these loci were identified. The normalized expression of these gene lists is shown for each disorder with alternating colors (black and blue). For each gene the developmental period with the maximum log_2_CPM value was identified and the mean period across the genes in each disorder is indicated with a point. **B)** The relative proportion (left, dashed lines represent expectation) and gene set enrichment (right) for trait/disorder-associated genes to the three trajectory groups (Figure 2). **C)** The analysis in ‘B’ is repeated for WGCNA modules. **D)** The analysis in ‘B’ is repeated for genes enriched for cell type clusters from scRNA-seq of the prenatal human brain. **E)** The analysis in ‘D’ is repeated for snRNA-seq of the adult human DLPFC. Abbreviations: DD: developmental delay; ASD: autism spectrum disorder; EPDD: epilepsy with developmental delay; GWAS: Genome-wide association study; ADHD: Attention deficit/hyperactivity disorder; MDD: major depressive disorder; EA: educational attainment; SCZ: schizophrenia; MS: multiple sclerosis; PD: Parkinson’s disease; AD: Alzheimer’s disease. Statistical analysis: B: FET, corrected for 30 comparisons. C: FET, corrected for 190 comparisons. D: FET, corrected for 190 comparisons. E: FET, corrected for 290 comparisons.

### Enrichment of CNS traits and disorders in co-expression and single cell gene lists

An alternative approach to identifying convergence across genes within a trait or disorder is to use co-expression (Gandal et al., 2018; Li et al., 2018; Parikshak et al., 2013; Willsey et al., 2013); therefore, we assessed enrichment for disease and trait-associated genes across the 19 WGCNA modules (Figure 3). Two modules are enriched after correcting for multiple comparisons (Figure 6C). The M4 Yellow module is enriched for genes associated with ASD and EA, including *NRXN1*, *TCF4*, and *BCL11A*, with a nominally significant trend in ADHD, DD, MDD, and SCZ. The M9 Brown module is enriched for genes associated with DD and EA, including *CDK13*, *PACS1*, and *EP300*, with a nominally significant trend in ASD and SCZ. Other modules in group 3 (M10 Dark-green, M11 Magenta, M12 Cyan) show nominally significant trends in multiple disorders and these include many of the genes with the strongest evidence for association with ASD and DD, including *ARID1B* in M10, and *ANKRD11*, *SETD5*, and *SYNGAP1* in M11.

Enrichment of cell type-specific genes in disorder-associated genes has been used to implicate cell types in disease pathology (Li et al., 2018; Satterstrom et al., 2018; Willsey et al., 2013; Xu et al., 2014). Using the gene lists described above (Figure 2), we perform a consistent analysis of cell type gene enrichment across the ten CNS traits and disorders (Li et al., 2018; Nowakowski et al., 2017). Five clusters showed significant enrichment, all from the fetal single cell data: C18 Excitatory newborn neurons and C1 Striatal interneurons in ASD, C3 Early excitatory neurons and C6 MGE-derived interneurons in EA, and C15 CGE-derived interneurons in EPDD (Figure 6D and 6E). Considering the nominally significant results too, we observe enrichment for a mixture of excitatory and inhibitory lineages in DD, ASD, EA, EPDD, and PD. For these disorders, the results in postnatal data are less pronounced, but not contradictory. For MS and AD, we observe nominally significant enrichment in prenatal microglia cluster-specific genes, while the postnatal data shows nominally significant enrichment for interneurons and astrocytes in MS and excitatory and inhibitory neurons in AD. Overall, protein-coding genes associated with a trait or disorder were more likely to be enriched in a cell type cluster in fetal single cell data (23.6% of trait or disorder genes vs. 16.1% of non-associated genes; OR=1.6 (95%CI:1.4-1.9); p=1.5×10^−10^ FET).

### eQTLs in CNS traits and disorders

We next considered whether the eGenes and eQTLs identified in this dataset overlapped with the genes and variants identified in the four GWAS analyses with the most loci: EA, SCZ, MS, and AD. To assess whether GWAS loci were enriched for eQTLs, as has been observed previously (Fromer et al., 2016; Nicolae et al., 2010), we used a permutation-based method that assessed expectation based on SNPs with similar LD structure, MAF, and gene density. Correcting for multiple comparisons, we find that SCZ, EA, and MS, but not AD, are enriched for eQTLs, with the lowest p-values obtained with Constant eQTLs rather than temporally specific eQTLs (Figure 7A), though we note that these temporal-specific enrichments are driven by small numbers of overlapping loci (Table S7). A complementary analysis, considering if eGenes are enriched for GWAS signal, did not show evidence of enrichment (Figure S6). Given that GWAS loci are enriched for eQTLs, we assessed whether the specific variants with strongest evidence at GWAS loci for EA and SCZ also had the strongest evidence of acting as eQTLs. This co-localization analysis showed that, while the loci overlapped, we did not see statistical evidence of overlap between the specific variants implicated in GWAS and acting as eQTLs (Figure S6).

**Figure 7.**
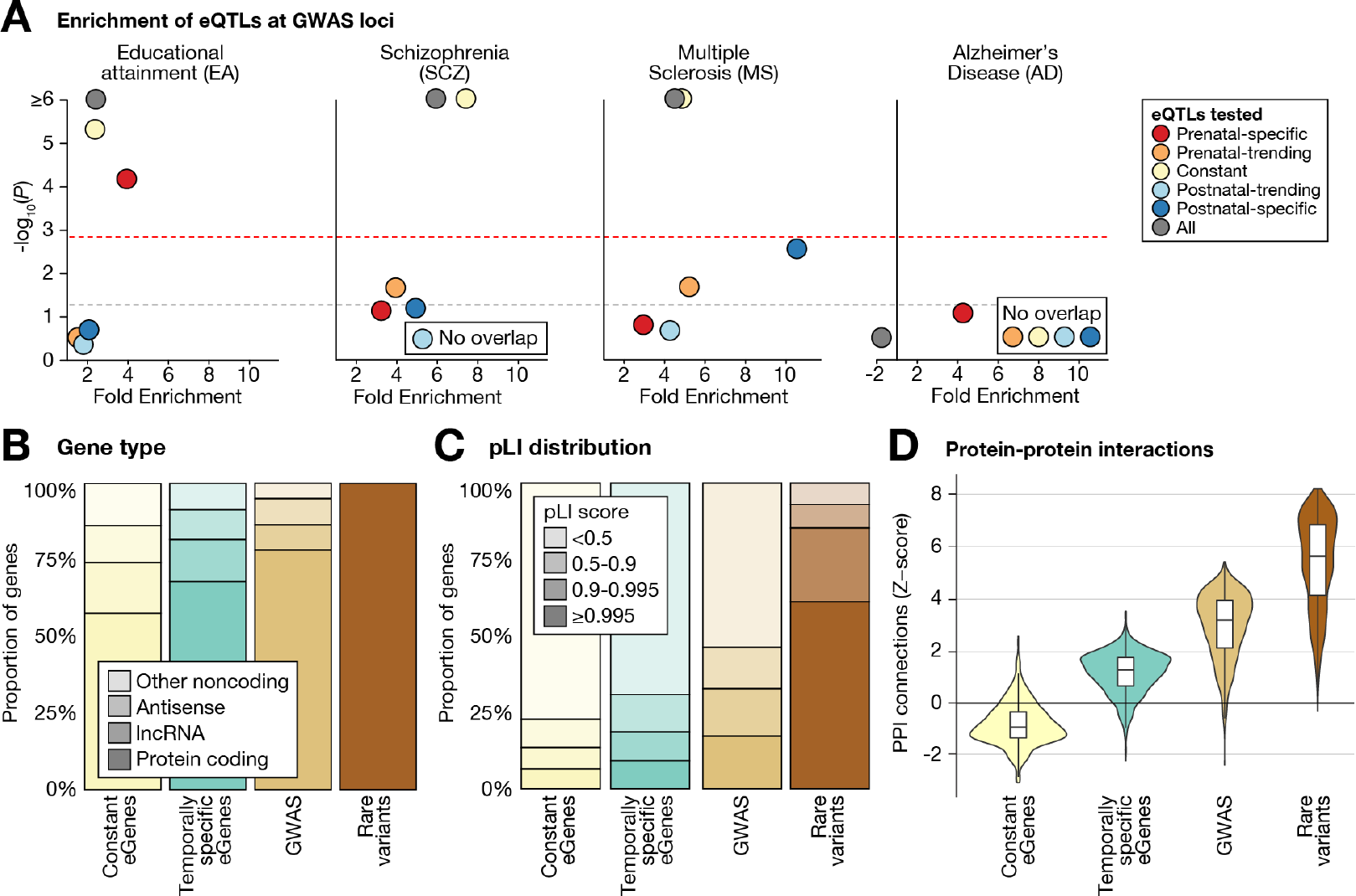
Relationship of eQTLs to genes associated with CNS traits and disorders. **A)** Enrichment of the eQTLs is shown for the four CNS GWAS datasets with p-value plotted against the degree of enrichment. Gray dashed line indicates nominal significance; red dashed line indicates significance threshold after correcting for 30 multiple comparisons. **B)** The relative proportions of coding and noncoding genes are shown for four gene sets defined by: eQTLs (Constant or Temporally specific, with Prenatal-specific and Postnatal-specific eGenes considered together, Figure 4), the union of the seven GWAS datasets, and the union of the three exome sequencing datasets (Rare variants). **C)** The distribution of pLI scores are shown for protein-coding genes in these four gene sets. **D)** BioGRID protein-protein interaction data was used to assess clustering on a network (Ripley’s K-net function), shown as a z-score, for these four gene sets. Abbreviations: GWAS, genome-wide association study; pLI, probability of loss-of-function intolerance. Statistical analysis: A: Permutation procedure comparing observed proportion of GWAS-significant SNPs that were eQTLs to a null distribution matched on LD structure, MAF, gene density and corrected for 30 comparisons.

To further understand the relationship between eQTLs and variants associated with human traits and disorders, we assessed the characteristics of the genes in four categories: Constant eQTLs, temporally specific eQTLs, GWAS loci (ADHD, EA, MDD, SCZ, MS, PD, AD), and Rare variant loci (DD, ASD, EPDD). The genes in proximity to GWAS loci are more likely to be protein-coding (Figure 7B), have high pLI scores (Figure 7C), and have multiple protein-protein interactions (Figure 7D) to a greater extent than temporally specific eQTLs (Figure 4) and similar to disorder-associated genes identified through rare variants.

## Discussion

In this manuscript, we describe BrainVar, a unique resource of paired genome (WGS) and transcriptome (bulk tissue RNA-seq) data, derived from 176 human dorsolateral prefrontal cortex samples across prenatal and postnatal development (Figure 1). We leverage these data to identify 23,782 genes expressed during human cortical development, gene lists relating to developmental trajectories and co-expression, common and rare variants that alter gene expression (eQTLs), and how these datasets relate to cell type, derived from scRNA-seq data, and CNS traits and disorders, derived from genomic analyses (exome sequencing and GWAS). In addition to developing a resource with utility for future studies on human development, neurobiology, and neuropsychiatric disorders, we also describe key biological insights, including the nature of the late-fetal transition in gene expression (Figure 2 and 3), the relative independence of variation in gene expression across development compared to gene expression between individuals (Figure 4), the variable penetrance of many haploinsufficient neuropsychiatric disorders (Figure 5), the developmental processes and cell types implicated in CNS traits and disorders (Figure 6), and the distinctive characteristics of genes with Constant eQTLs compared to those with temporally specific eQTLs or associated with CNS traits and disorders (Figure 4 and 7). Across these analyses, we observe complicated relationships between gene function and development, consistent with the impact of natural selection.

Principal component analysis identifies developmental age as the most important factor underlying the variance in gene expression in this dataset. The majority of this temporal variance occurs in two transitional phases (Figure 2). The early-fetal transition is a coordinated decrease in expression of multiple genes in early development (epoch 0; periods 1-2; 6-10 PCW), captured in the lncRNA-enriched M8 Red module (Figure 3). This transition may relate to the establishment of regional identity across the brain since it includes genes associated with identity of non-cortical tissues, however the paucity of samples in this stage limits our interpretation of this shift. The late-fetal transition between mid-fetal development and infancy involves over 12,000 genes, with similar numbers rising and falling (Figure 2). Prior reports in humans (Li et al., 2018) and primates (Zhu et al., 2018) associated this transition with a reduction in intra- and inter-regional variation evident at the levels of bulk tissue and individual cell types. In keeping with these initial reports, our data suggest this transition represents a combination of changes in the relative proportions of various cell types and biological processes within these cells (Figure 2 and 3). Specifically, the late-fetal transition appears to be the co-occurrence of decreasing cell division (M2 Royal-blue), neuronal differentiation (M4 Yellow) and anabolic processes (M1 Black) with increasing gliogenesis (M18 Light-cyan, M19 Turquoise), neuronal maturation and synaptogenesis (M19 Turquoise), and the metabolic demands of supporting neuronal activity (M13 Green, M14 Purple, M16 Blue). Critically, this larger sample set allowed us to better define the inflection point at which the late-fetal transition begins as being within period 5, further distinguishing the late-fetal transition from previously reported organotypic changes (Domazet-Loso and Tautz, 2010; Kalinka et al., 2010; Li et al., 2018).

While previous analyses have identified eQTLs in human brain tissue both postnatally, (Fromer et al., 2016; The GTEx Consortium, 2015) and prenatally (O’Brien et al., 2018), this dataset has two unique characteristics. First, no prior study has assessed the impact of genomic variation in the context of gene expression across the whole of brain development, from embryogenesis through fetal development, infancy, childhood, adolescence, and into young adulthood.

Consequently, we were able to identify temporally specific eQTLs that only modulate expression prenatally or postnatally. Second, by using deep coverage WGS rather than genotyping arrays to detect various types of genomic variation, we were able to assess the impact of both common and rare variants on gene expression.

For common variation, we identify 252,629 eQTLs that act on 8,241 of the 23,782 cortically-expressed genes. The majority are Constant eQTLs, observed in both the prenatal and postnatal samples, while 91,336 eQTLs (36.1%) are temporally specific and act on 5,444 genes (Figure 4A). Considering the FDR-significant variant-gene pairs that overlap with prior eQTL catalogues (O’Brien et al., 2018; The GTEx Consortium et al., 2017) we observe high concordance in both the direction and magnitude of effect (Figure 4C), despite differing in cohorts, methods, and analysis. Concordance was greatest for Constant eQTLs and temporally specific eQTLs from cohorts of equivalent developmental stages (Figure 4C).

Across multiple metrics, we observe dramatic differences between the eGenes with Constant and temporally specific eQTLs (Figure 4 and 7). Compared to other genes expressed in the cortex, the genes affected by Constant eQTLs are more likely to be noncoding, be weakly co-expressed, and have few protein-protein interactions. In contrast, the genes regulated by eQTLs with a degree of temporal specificity are similar to genes for which we did not detect eQTLs. A similar distinction was observed for distance from the TSS, with Constant eQTLs close to the gene (~90kb), while temporally specific eQTLs are further (~420kb). Critically, we find that pLI score, a measure of sensitivity to variation in gene expression, and eQTL effect size are inversely related (Figure 4K). Furthermore, Prenatal-specific eGenes are more common among Rising genes with low prenatal expression. These observations suggest that developmental and evolutionary constraints limit the population frequency or impact of eQTLs on key developmental processes, a hypothesis that might be testable in future studies as additional information concerning spatiotemporal and cell type specificity of enhancers and eQTLs becomes available for a variety of tissues. Under this model, Constant eQTLs with high effect sizes tend to influence genes that tolerate variation in expression (e.g. non-rate-limiting metabolic steps) or are non-critical to brain function, while temporally specific eQTLs tend to influence genes with critical roles that are sensitive to variation in expression, but only to a small degree or at a stage in development when expression variation is tolerated. These temporally specific effects appear to be modulated through distal regulatory elements, perhaps reflecting the influence of transcription factors with temporal and/or cell type specificity.

Relationships between developmental stage, cell type specificity and other confounding factors, such as natural selection, are evident in other aspects of our dataset. For example, although GWAS loci in SCZ, MS, and EA are enriched for eQTLs (Figure 7A), colocalization analysis suggests that different variants mediate the effects in each dataset (Figure S7). A potential explanation may lie in the observation that those characteristics distinguishing temporally specific eQTLs from Constant eQTLs, such as higher pLI scores and interactions with more proteins, are exaggerated at GWAS loci and disorder-associated rare variant loci. Alternatively, neuropsychiatric disorders may be mediated by cell types and subtypes, which our study is not powered to detect. Indeed, relationships between genes associated with neuropsychiatric disorders and specific cell types are evident in the data presented in this manuscript, as we observe the enrichment of genes associated with DD, ASD, and EA in developing neurons of the excitatory and inhibitory lineages in the prenatal brain and increased associations between late-onset disorders and glial cell types (Figure 6). eQTLs derived from single cell expression data or other brain regions may provide greater clarity. Also suggesting differing specificity or other confounding issues, the magnitude of effects we observed for rare heterozygous PTVs and deletions was less than anticipated and inconsistent between variants, potentially reflecting a dilution of cell type specific signals against a background of inter-individual variation, which may explain variable penetrance, or other effectors including autoregulation, protective alleles, alternate isoforms, or an incomplete model of PTV effects (MacArthur et al., 2012; Rivas et al., 2015). Higher resolution datasets across development, including single cell, additional brain regions, and larger sample sizes, along with complementary analyses in the brains of individuals with neuropsychiatric disorders and rare genetic disorders, are likely to provide additional insights into these relationships.

The combination of genomic and transcriptomic data across development allows us to interrogate human cortical development from a molecular perspective at a higher resolution than before. Understanding patterns of temporal and cell type specificity has already provided insights into the pathology underlying neuropsychiatric disorders and further delineation of these patterns is likely to be critical to a detailed understanding of etiology as a foundation for therapeutic development.

## Supporting information

Table S1 Samples

Table S2 Genes

Table S3 WGCNA

Table S4 eQTLs

Table S6 PTVs and deletions

Table S6 Disorders

Table S7 GWAS overlap

## Acknowledgments

Data were generated as part of the PsychENCODE Consortium, supported by: U01MH103339, U01MH103365, U01MH103392, U01MH103340, U01MH103346, R01MH105472, R01MH094714, R01MH105898, R21MH102791, R21MH105881, R21MH103877, and P50MH106934 awarded to: Schahram Akbarian (Icahn School of Medicine at Mount Sinai), Gregory Crawford (Duke), Stella Dracheva (Icahn School of Medicine at Mount Sinai), Peggy Farnham (USC), Mark Gerstein (Yale), Daniel Geschwind (UCLA), Thomas M. Hyde (LIBD), Andrew Jaffe (LIBD), James A. Knowles (USC), Chunyu Liu (UIC), Dalila Pinto (Icahn School of Medicine at Mount Sinai), Nenad Sestan (Yale), Pamela Sklar (Icahn School of Medicine at Mount Sinai), Matthew State (UCSF), Patrick Sullivan (UNC), Flora Vaccarino (Yale), Sherman Weissman (Yale), Kevin White (UChicago) and Peter Zandi (JHU). This work was supported by funding provided by the Autism Science Foundation (DW, JA, SJS); Simons Foundation Autism Research Initiative (SFARI) grants #574598 (SJS) and #402281 (SJS, MWS, BD, KR); National Institute for Mental Health (NIMH) grants R01 MH109901 (SJS, MWS), R01 MH110928 (SJS, MWS), U01 MH103339 (MWS), R01 MH111662 (SJS, MWS), U01 MH105575 (MWS), U01 MH106874 (NS), P50 MH106934 (NS), R01 MH109904 (NS), R01 MH110926 (NS), U01 MH116488 (NS); and National Institute of Child Health and Human Development (NICHD) grant R24 HD000836 (IAG). We thank Thomas Lehner, Anjené Addington, Geetha Senthil, and Alexander Arguello at the NIMH for their support of the PsychENCODE consortium, the Yale Center for Genome Analysis for generating the RNA-seq data, GENEWIZ for generating the WGS data, and Sentieon for the use of a computationally efficient implementation of GATK Haplotype Caller.

## Author Contributions

Conceptualization, KR, MWS, BD, SJS, and NS; Methodology, DMW, SP, JC, JA, HAB, JLT, KAA, DRO, IAG, KR, MWS, BD, SJS, and NS; Software, DMW, JA, CD, LL, MCG, and SJS; Validation, DMW, SP, JC, JA, CD, SD, BD, SJS, and NS; Formal Analysis, DMW, SP, JC, JA, BS, MP, GS, MSB, LK, AEC, BD, and SJS; Investigation, DMW, SP, JC, JA, BS, BD, SJS, and NS; Resources, SP, JC, ZL, CD, AMS, ATT, NK, FOG, LL, MCG, XZ, HAB, JLT, KAA, DRO, IAG, MET, SJS, and NS; Data Curation, DMW, SP, JC, CD, LL, MCG, XZ, SD, and SJS; Statistical analysis, DMW, JC, JA, BS, MP, MSB, BD, and SJS; Writing Original Draft, DMW, SP, JC, JA, BS, BD, SJS, and NS; Writing Review & Editing, DMW, SP, JC, JA, BS, MP, ZL, CD, GS, AMS, ATT, NK, FOG, MSB, LL, MCG, XZ, SD, LK, AEC, JDB, HAB, JLT, KAA, DRO, IAG, NAZ, MET, KR, MWS, BD, SJS, and NS; Visualization, DMW, SP, JC, JA, BS, SJS, and NS; Supervision, JDB, NAZ, MET, KR, MWS, BD, SJS, and NS; Project Administration, DMW, SP, SJS, and NS; Funding Acquisition, KR, MWS, BD, SJS, and NS.

**Declaration of Interests**

The authors declare no competing interests

