## Supplementary Methods for "Whole-genome and RNA sequencing reveal variation and transcriptomic coordination in the developing human prefrontal cortex"

#### CONTACT FOR REAGENT AND RESOURCE SHARING

#### EXPERIMENTAL MODEL AND SUBJECT DETAILS

##### Human postmortem tissue

This study was conducted using postmortem human brain specimens from tissue collections at the Department of Neuroscience at Yale University School of Medicine. Additional specimens were procured from the Birth Defects Research Laboratory at the University of Washington, Biobank BB-0033-00064, Department of Pathology, Lariboisière Hospital (Paris), Advanced Bioscience Resources Inc., Human Brain Collection Core (HBCC), the Brain and Tissue Bank at the University of Maryland, the MRC-Wellcome Trust Human Developmental Biology Resource at the Institute of Human Genetics, University of Newcastle, UK, the Human Fetal Tissue Repository at the Albert Einstein College of Medicine (AECOM), the Human Brain Collection Core (HBCC), and the Brain and Tissue Bank at the University of Maryland. Tissue was collected after obtaining parental or next of kin consent and with approval by the institutional review boards at the Yale University School of Medicine, the National Institutes of Health, and at each institution from which tissue specimens were obtained. Tissue was handled in accordance with ethical guidelines and regulations for the research use of human brain tissue set forth by the NIH (http://bioethics.od.nih.gov/humantissue.html) and the WMA Declaration of Helsinki (http://www.wma.net/en/30publications/10policies/b3/index.html).

All available non-identifying information was recorded for each specimen in Table S1. In total, 176 postmortem brain specimens (104 male, 72 female; postmortem interval of 21.7 ± 15.9 (mean ± SD) hours and pH, 6.41 ± 0.35) ranging in age from 6 post-conception weeks (PCW) to 20 postnatal years (PY) (Figure 1 and Table S1) were included in this study. Fetal age was extrapolated based on the date of the mother's last menstruation, characteristics of the fetus noted upon ultrasonography scanning, foot length of the fetus, and visual inspection. The postmortem interval (PMI) was defined as hours between time of death and time when tissue samples were frozen.

#### METHOD DETAILS

##### Tissue dissection

Tissue was dissected as described previously (Kang et al. 2011). Samples collected from 6 – 9 PCW specimens contained the entire thickness of the cerebral wall. Samples collected from 12 – 22 PCW specimens contained the cortical plate. Samples from 35 PCW – 20 PY specimens were dissected such that the entire gray matter (layer 1-6) and part of the underlying subplate (4 – 12 postnatal months) or white matter (1 – 20 PY) were collected.

##### RNA extraction and quality assessment

Total RNA was extracted using mirVana kit (Ambion) with some modifications, as given in the following text, to the manufacturer's protocol, as described below. Each tissue sample was pulverized with liquid nitrogen in a prechilled mortar and pestle and transferred to a chilled safe-lock microcentrifuge tube (Eppendorf). Per tissue mass, equal mass of chilled stainless steel beads (Next Advance, catalog # SSB14B) along with one volume of lysis/binding buffer were added. Tissue was homogenized for 1 min in Bullet Blender (Next Advance) and incubated at 37°C for 1 min. Another nine volumes of the lysis/binding buffer were added, homogenized for 1 min, and incubated at 37°C for 2 min. One-tenth volume of miRNA Homogenate Additive was added and extraction was carried out according to the manufacturer's protocol. RNA was treated with DNase using TURBO DNA-free Kit (Ambion/Life Technologies) and RNA integrity was measured using Agilent 2200 TapeStation System.

**RNA-seq library preparation and sequencing**
Barcoded libraries for RNA-seq were prepared with 5ng of RNA using TruSeq Stranded Total RNA HT Sample Prep Kit with Ribo-Zero Gold kit (Illumina) per manufacturer's protocol. Paired-end sequencing (100 bp x 2) was performed on HiSeq 4000 sequencers (Illumina) at Yale Center for Genome Analysis.

**DNA extraction**
Genomic DNA was isolated using the QIAamp DNA Mini Kit (Qiagen). In detail, approximately 25 mg of brain tissue was transferred to a chilled safe-lock microcentrifuge tube (Eppendorf) and equal mass of chilled stainless-steel beads (Next Advance, catalog # SSB14B) along with 90 µl of buffer ATL were added. Tissue was homogenized for 1 min in Bullet Blender (Next Advance) and incubated at 37°C for 1 min. Another 90 ul of buffer ATL was added and blended for an additional minute. After incubation on ice for 5 min, tubes were gently centrifuged to collect beads at the bottom. Supernatant was transferred to a new tube and 20 µl of Proteinase K was added. Sample was incubated at 56° C for 3 hours in a shaking heat block. After incubation, genomic DNA was further purified following the manufacturer's protocol. DNA was eluted in nuclease free water and concentration was estimated by nanodrop.

**Whole-genome sequencing**
DNA library preparation and sequencing were carried out at GENEWIZ (New Jersey). Before library preparation, the concentration of the DNA was measured using a fluorescent assay and DNA quality was assessed by visualization on agarose gels. PCR-free DNA library preparation was performed and resulting libraries were sequenced at 2x150 bp to achieve mean coverage of 30x (Table S1).

### QUANTIFICATION AND STATISTICAL ANALYSIS

**WGS variant calling**
Using the pipeline from the Centers for the Common Disease Genomics project (Regier et al., 2018), FASTQ reads were aligned to the GRCh38 reference from the 1000 Genomes Project using BWA-MEM version 0.7.15. Reads were sorted and duplicates were removed with Picard, version 2.17.5.; base quality score recalibration was then performed with the Genome Analysis Toolkit (GATK), v3.8-0-ge9d806836. Variant calling and joint genotyping were done with the Haplotyper and Genotyper tools from Sentieon v201711.01**,** a toolkit containing modules that are mathematically equivalent to their counterparts in the GATK (Freed et al., 2017). SNP and indel recalibration were performed on the joint genotyped VCF file. Variant Quality Score

Recalibration (VQSR) metrics were created from a training set of highly validated variant resources: dbSNP build 138, HapMap 3.3, 1000 Genomes OMNI 2.5, and 1000 Genomes Phase 1. For the following analyses, we excluded: variant calls with any VQSR tranches (keeping "PASS" only), variants located in low-complexity regions (Li, 2014), variants located on non-canonical chromosomes (decoy chromosomes or contigs), indels >50 bp, SNVs with allele balance >0.78 or <0.22 (indels >0.8 or <0.2), variants with <90% call rate, and variants and genotypes that did not meet high quality thresholds as identified in an ROC-based optimization procedure using family-based WGS data (Werling et al., 2018).

Variants with a minor allele frequency of ≥5% in both the prenatal (periods 1-6; N=112) and postnatal (periods 8-12; N=60) samples were included in downstream expression quantitative trait locus (eQTL) analysis (N=6,573,196 variants). For annotation and subsequent analyses, we converted the final VCF into Variant Dataset Format using Hail version 0.1. SNPs and insertions/deletions (up to 50bp) annotation based on the GENCODE comprehensive version 21 (Harrow et al., 2012) using Ensemble VEP version 90 (McLaren et al., 2016).

**RNA-seq alignment and gene-level read count quantification**
RNA-seq reads were aligned to the human genome (hg38/GRCh38) using STAR aligner (Dobin et al., 2013) and gene-level read counts were calculated using HTSeq (Anders et al., 2015) based on GENCODE v21 annotation (Harrow et al., 2012).

**RNA-seq normalization and technical artifact correction**
The read count data matrix of 176 samples by 60,155 genes (Table S2) was normalized as follows:

- Step 1: Read count data matrix converted to counts per million (CPM).
- Step 2: Genes with CPM ≥1 CPM in at least 50% of the samples in any one sex in any one period were included; 23,782 genes passed these criteria.
- Step 3: CPM values were transformed to $\log_2$(CPM) using the voom function in the limma R package (Law et al., 2014; Ritchie et al., 2015).
- Step 4A: In order to correct for the technical artifacts, we performed hidden covariate analysis on the residuals of the expression matrix after developmental period and sex were subtracted from the $\log_2$(CPM) data matrix using the hidden covariate analysis method (HCP) (Mostafavi et al., 2013).
- Step 4B: In parallel, we performed surrogate variable analysis (SVA) on residuals of the expression matrix after developmental period and sex were subtracted from the $\log_2$(CPM) data matrix using the SVA R package (Leek et al., 2012).
- Step 5: In each of Step 4A and 4B, we subtracted contributions from 20 hidden covariates (HCP) and 2 surrogate variables (SVA) from the $\log_2$(CPM) data matrix.

Unadjusted $\log_2$CPM gene expression data for 176 samples by 23,782 genes was used for expression trajectory (Figure 2) and WGCNA analyses (Figure 3). While the adjusted (HCP/SVA) values were used for eQTL analysis.

**Data quality and sample identity assessment**

To confirm that the WGS and RNA-seq data from each sample were of sufficient quality for downstream analysis and corresponded to the same individual, a series of quality metrics and checks were performed (Table S1).

For the WGS data, coverage metrics were assessed using PicardTools (v2.18.1). Mean coverage per sample ranged from 22.7-65.4x, with a cohort median of 31.5x. Across all samples, 92.1-93.6% of the mapped genome was covered at 10x or greater (mean of 92.8%). The FREEMIX metric from VerifyBamId (v1.1.3; (Jun et al., 2012)) was used to identify samples with potential contamination, with a maximum observed FREEMIX score of 0.064, suggesting no contamination (Lek et al., 2016).

Sample identity was verified by comparing sex and genotype between the WGS and RNA-seq data. In the WGS data, sex was determined from chromosome X heterozygosity using Peddy (v0.3.2; (Pedersen and Quinlan, 2017)), with the Peddy hg19.sites converted to GRCh38 using the UCSC Genome Browser LiftOver utility. High-quality variants with an allele frequency ≥1% were exported from the VCF using Hail for input into Peddy. In the RNA-seq data, sex was determined from the expression levels of *XIST* and the 18 most highly expressed genes on chromosome Y: *KDM5D*, *DDX3Y*, *ZFY*, *TBL1Y*, *PCDH11Y*, *PRKY*, *USP9Y*, *RPS4Y1*, *TXLNGY*, *NLGN4Y*, *TTTY14*, *UTY*, *EIF1AY*, *GYG2P1*, *TTTY10*, *TTTY15*, *KALP*. Based on gene-specific expression thresholds determined by visual inspection of bi-modal expression histograms, each sample's sex was predicted according to the expression level of all 19 genes. Sex was consistent in the WGS and RNA-seq data for all 176 samples and matched the recorded sex in 132 out of 134 samples with such data (55/56 females, 77/78 males).

To confirm identity by genotype, we compared the genotypes from 289 common, coding SNPs with high fixation index (FST) (Sanders et al., 2015), called from both the WGS and RNA-seq data. Genotypes were callable in both data types for 118-206 SNPs per sample (40.8-71.3% of 289 SNPs; median = 177, 61.2%). SNP variant genotypes were highly concordant between the WGS and RNA-seq data (median 87.4% concordance between WGS and RNA-seq for the corresponding sample; lowest concordance 73%), with corresponding samples showing higher concordance than comparisons between all discordant samples. There was no evidence of duplicate or closely related samples (SNP-based relatedness coefficients from Peddy: -0.000332 to 0.1481).

To confirm the approximate accuracy of samples' reported age, the expression level of the doublecortin gene (*DCX*) was examined. DCX is involved in neuron migration, and is expressed most strongly during prenatal development, with distinctly decreased postnatal expression. All 176 samples showed the expected DCX expression levels given samples' reported age. Similar results across all expressed genes were obtained using principal component analysis (below).

**Ancestry estimation**

Ancestry was estimated using principal component analysis of common SNPs and indels in the WGS data, run alongside 3,804 additional individuals of known ancestry with WGS data (parents from the Simons Simplex Collection (An et al., 2018)). From 10,688,106 SNPs and indels with allele frequency ≥5% in either this data set, one of the three batches of Simons Simplex Collection data, or GnomAD genomes, variants were pruned for independence with linkage disequilibrium $r^2$<0.1 and then randomly downsampled to 118,849 variants. Principal component analysis was run using Hail 0.1. The first two principal components were used to classify samples by ancestry, and the first five principal components were used as covariates in the identification of eQTL loci (Figure S1).

**Estimation of biological and technical covariates in RNA-seq data**
PCA analysis was performed on the covariance matrix of 23,782 cortically-expressed genes in 176 samples. A secondary PCA was performed on the 11,705 Non-transitional genes in 167 samples (excluding Period 1 and 2 samples) to assess the extent to which removing the late-fetal transition accounted for variance in gene expression. For each PCA, the variance explained by each principal component was assessed (Figure S2).

To quantify the relative contributions of biological and technical covariates, we calculated the partial $R^2$ of each covariate with each principal components using the rsq R package, in a generalized linear model where loadings of principal components are considered as a response and biological and technical covariates (such as developmental period, sex, sequencing batch, sequencing depth, RNA integrity number (RIN), mitochondrial RNA proportion, ribosomal RNA proportion, intronic reads proportion, intergenic reads proportion) are considered as predictor variables (Figure S2).

**Transcriptome temporal trajectory estimation**
Statistically, the temporal dynamics of the expression of genes can be modeled as a mixture of K distinct trajectories, each with Gaussian noise (Jones et al., 2001; Roeder et al., 1999). To delineate the temporal dynamics for K groups of genes, we used the Flexmix R package, which provides the expected trajectory for each group and the soft group assignments of individual genes to groups.

The expression of 23,782 genes was transformed as $\log_2$(CPM) and normalized by the interquartile range. The samples in epoch 0 were excluded to avoid biased estimation due to very few samples. To identify the overall trend of expression over age, first we fitted the model on all the 23,782 genes assuming there are three groups and that the expected trajectories for each group can be represented with degree-4 polynomials on age. Three typical trajectories were identified, including a group of 6,941 genes with Rising expression levels, a group of 5,173 with Falling expression levels, and a group of 11,705 genes with roughly flat (fitted) expression over time, which we called Non-transitional (Figure 2A).

**Gene ontology functional enrichment for temporal trajectories**
For functional enrichment, we characterized genes sets for each trajectory using the R package, gProfiler (Reimand et al., 2011). The pathway enrichment test was performed using Gene Ontology Biological Process terms, which contain between 10 and 2,000 genes, and all 23,782 cortically-expressed genes were used as background. Enrichment tests were subject to the “moderate” hierarchical filtering parameter, and the FDR multiple correction in the gProfiler (Figure 2).

**Assessing enrichment in tissue-specific genes from GTEx**

To assess enrichment across tissues, 27,546 transcripts with an RPKM ≥0.5 in 80% of samples one or more tissues in GTEx (gtexportal.org) were defined and log-transformed ($log_2$[RPKM+1]). For each gene, expression between each tissue and all other tissues was assessed using a moderated t-test (R package limma), with models adjusted for age, RIN, gender, and surrogate variables. The Benjamini and Hochberg method was used to estimate false discovery rate (FDR) and tissue enriched genes were defined as: log fold-change > 0.5 and FDR<0.05. The enrichment of the Falling, Non-transitional, and Rising genes was assessed with using Fisher's exact test with 23,782 cortically-expressed genes as a background (Figure 2).

**Identifying genes enriched in cell types from single cell data**

To identify cell type enriched genes we calculated a tau metric (Kryuchkova-Mostacci and Robinson-Rechavi, 2017) from the per gene $log_2$TPM of $log_2$UMI values for genes within cell type clusters in prenatal forebrain (Nowakowski et al., 2017) and postnatal cortex (Li et al., 2018). Each gene was also ranked across all clusters within the dataset on the basis of TPM/UMI, so that the cluster with the highest TPM/UMI was ranked as "1", while the cluster with the second highest TPM/UMI for that gene was ranked as "2", etc. The genes were sorted by TPM/UMI rank, tau, and absolute TPM.UMI. The top 200 genes from this list were selected so long as the TPM/UMI rank was ≤12, the tau was ≥0.3, and the absolute TPM/UMI was ≥-3. For the prenatal single cell genes, the mean TPM rank was 2.48, the mean tau was 0.68, and the mean absolute TPM was 0.50. For the postnatal data the mean UMI rank was 1.09, the mean tau was 0.76, and the mean absolute UMI was 0.31.

**Enrichment of gene trajectories in temporal putative cis-regulatory elements**

H3K27ac peaks present in more than two samples of fetal or adult dorsal frontal cortex in BrainSpan (Li et al., 2018) were tested for fetal versus adult temporal bias using DESeq2 (Love et al., 2014). Temporally biased genes were defined as adjusted P<0.01 and fold change ≥2. A category of "non-temporal" H3K27ac peaks was generated with peaks showing P>0.05. All peaks were annotated using the gene with the closest transcription start site in Gencode v21. Genes were classified as only-fetal or only-adult if they were associated with fetal or adult-only H3K27ac peaks, respectively. Enrichments in of each category of H3K27ac-genes in each category of eGenes were tested by means of a Fisher Exact's test and P-values were adjusted using Benjamini-Hochberg, using genes associated to non-variant H3K27ac peaks as a reference background.

**WGCNA network construction and module definition**

To assess the functional topology in cortical samples, we applied Weighted Gene Co-Expression Network Analysis (WGCNA) (Langfelder and Horvath, 2008) to 23,787 cortically-expressed transcripts. Network analysis was performed with WGCNA (version 1.63) using a signed network, choosing a soft-threshold power, the mean connectivity less than 50, and scale-free topology greater than 0.8. To reduce the bias driven by a few sample outliers, we applied the blockwiseConsensusModules function, which detects consensus modules across 100 subsampled networks. We used the average linkage hierarchical clustering of the topological overlap dissimilarity matrix (1-TOM) to generate the network dendrogram. Modules were defined as branches of the dendrogram using the hybrid adaptive tree cut with the following parameters: minimum module size = 200, negative pamStage, height cut = 0.999, and deep split = 2 (Langfelder and Horvath, 2007). Modules were summarized by their first principal component

(ME, module eigengene), followed by merging modules with high correlations (eigengene value ≥0.9).

**WGCNA functional enrichment for module characterization**

For functional enrichment, we characterized WGCNA module genes using the gProfiler R package (Reimand et al., 2011), as described above in the analysis of temporal trajectory genes. To identify WGCNA module genes that are regulatory targets, we searched for transcription factor binding targets using the ChEA (Lachmann et al., 2010; Satterstrom et al., 2018), and TRANSFAC (Matys et al., 2003), and microRNA using the mirTarbase database (Chou et al., 2018).

**WGCNA module preservation**

To assess whether 19 co-expression modules in our samples were preserved in other, independent DLPFC or frontal cortex expression datasets, we compared our dataset with non-overlapping samples from the BrainSpan dataset (Li et al., 2018) and applied the module preservation function from the WGCNA R package (Langfelder et al., 2011). From the BrainSpan dataset, we selected DLPFC (n=30), 10 non-DLPFC neocortical regions (n=317), and subcortical regions excluding the cerebellum (n=140). Given the original co-expression network constructed above, we used modulePreservation to calculate module preservation statistics from 100 permutations (Table S3).

**Clustering analysis in protein-protein interaction network**

To examine functional association of a group of genes/proteins, we performed a clustering analysis of a protein-protein interaction network implemented in SANTA R package (Cornish and Markowetz, 2014). Ripley's K function provides a measure of whether points are clustered together or randomly dispersed (homogenous) in a network and the SANTA R package reformulated Ripley's K function for a protein-protein interaction network. Clustering can be indicative of functional association between the genes/proteins under consideration. To test departure from homogeneity of a given gene set, we drew an empirical null distribution of clustering from 1,000 random samples of matching sized gene sets from the BioGRID protein-protein interaction network data (Winter et al., 2011) (v3.4.132). We reported departure from null distribution as a Z score (i.e., $Z > 0$: level of clustering of a gene set greater than null expectation and $Z < 0$: level of clustering of a gene set less than null expectation).

**Cis-eQTL detection and classification**

Cis-eQTLs were identified for all high quality, common variants (N=6,573,196) within 1 Mb of a gene boundary using the linreg function in Hail 0.1, with period, sex, and the first five principal components of common variant ancestry as covariates. This analysis was run on three cuts of the BrainVar data set: complete sample (N=176, periods 1-12), prenatal-only (N=112, periods 1-6), and postnatal-only (N=60, periods 8-12). Separately for the results of each analysis, false discovery rate (FDR) was calculated for all gene-variant pairs using the Benjamini-Hochberg procedure.

We then classified all gene-variant pairs with FDR≤0.05 from at least one analysis into groups defined by the temporal specificity of their eQTL effects. To do this, we first identified one variant per gene with the smallest, FDR-significant p-value, from any of the three analyses. We then used a Z-test to compare the regression coefficients for these variant-gene pairs from the prenatal and postnatal analyses:

$$Z = \frac{\beta_{\mathrm{Pre}} - \beta_{Post}}{\sqrt{SE_{\mathrm{Pre}}^2 + SE_{Post}^2}}$$

Using the results from the eQTL analyses and from this prenatal-postnatal comparison, we then classified each of these top (one per gene) gene-variant pairs and their corresponding target gene (eGene) into one of five groups:

1. Constant eQTLs/eGenes, characterized by consistent effects across this developmental data set: FDR≤0.05 in the complete sample analysis, same direction of effect and unadjusted p≤0.05 in both the prenatal and postnatal analyses
2. Prenatal-specific eQTLs/eGenes, with strongest effects during prenatal development: FDR≤0.05 in the prenatal analysis, unadjusted p>0.05 in the postnatal analysis, pre-post comparison Z-test FDR-adjusted p≤0.05
3. Postnatal-specific eQTLs/eGenes, with strongest effects during postnatal development: FDR≤0.05 in the postnatal analysis, unadjusted p>0.05 in the prenatal analysis, pre-post comparison Z-test FDR-adjusted p≤0.05
4. Prenatal-trending eQTLs/eGenes, which did not fit into earlier categories, but had higher prenatal effects ($B_{Pre} > B_{Post}$)
5. Postnatal-trending eQTLs/eGenes, which did not fit into earlier categories, but had higher prenatal effects ($B_{Post} > B_{Pre}$)

All FDR-significant variants associated with the expression of a single gene were classified into one of these five groups according to the classification of the top variant for the same gene (Figure 4).

**Alternative approaches for assigning eGenes to temporal categories**
Many eGenes are associated with multiple eQTLs, each of which could individually meet criteria for any one of the five temporal categories. As described above, we categorized eGenes into temporal categories based on the performance of their top eQTL (smallest p-value), but we assessed the performance against two alternative approaches: (1) eGene assigned to the same category as a majority of their eQTLs ("majority eQTL" approach), with ties assigned in the order Constant, Prenatal-specific, Postnatal-specific, Prenatal-trending, Postnatal-trending, and (2) for eGenes with ≥1 eQTL, category assignment based on the performance of the second most significant variant ("second eQTL" approach). For each of these alternative approaches, we calculated the percent of eGenes from each top variant-based category that were assigned to each category using the majority eQTL or the second eQTL approach (Figure S4).

**Comparison with published eQTL studies**
To assess the sensitivity of our cis-eQTL discovery analysis relative to previous work, we evaluated the relationship between sample size and eGene discovery for: the BrainVar prenatal, postnatal, and complete sample analyses as run using the HCP- and SVA-adjusted expression data, GTEx v7 analyses by tissue (gtexportal.org), postnatal human frontal cortex by the CommonMind Consortium (Fromer et al., 2016), and prenatal human whole brain (O'Brien et al., 2018). Using the sample size reported in each analysis, or for each tissue (GTEx), and the

number of genes with at least one eQTL reaching significance of FDR ≤ 0.05, we plotted the relationship between eGene discovery and sample size (Figure S4). We observe a strongly positive relationship across BrainVar and the published analyses, in keeping with prior reports that eQTL and eGene discovery is positively associated with sample size (The GTEx Consortium et al., 2017).

We evaluated the performance of the eQTLs that we identified in our analyses with published sets of eQTLs identified in the human postnatal frontal cortex (The GTEx Consortium et al., 2017) and in the human prenatal whole brain (O'Brien et al., 2018). For the postnatal frontal cortex data, we downloaded significant variant-gene pairs from the GTEx v7 data release from gtexportal.org and used R to write out the variant locations to a bed file format. We then used the command line LiftOver utility from the UCSC Genome Browser to convert the hg19 variant positions to GRCh38. For the prenatal brain data, we downloaded the eQTL summary statistics, results for the top eQTLs per gene, and SNP positions bed file from the study data repository on Figshare (https://figshare.com/articles/Summary_statistics_for_expression_quantitative_trait_loci_in_the_developing_human_brain_and_their_enrichment_in_neuropsychiatric_disorders/6881825). We then used each gene's nominal significance threshold from the top eQTLs file to identify the full set of variant-gene pairs meeting significance.

Using variant (GRCh38 position, reference, and alternate alleles) and gene (Ensembl gene IDs) identifiers, we matched significant variant-gene pairs separately from GTEx frontal cortex and prenatal brain to the variant-gene pairs meeting FDR significance (≤ 0.05) in the BrainVar analyses. For all significant eQTLs in BrainVar and for the eQTLs in each temporal category (Constant, Prenatal-specific, Postnatal-specific, Prenatal-trending, Postnatal-trending), we then calculated the number of eQTLs that were identified in both the BrainVar and reference analysis, the number of eQTLs unique to BrainVar, and the percentage of overlapping out of total BrainVar eQTLs. For eQTLs that overlapped, we then compared the effect of the variant on the expression of its associated gene to determine the percentage of overlapping eQTLs with concordant direction of effect, as well as the Pearson correlation between the eQTL effects (beta from BrainVar, slope from GTEx or prenatal whole brain). The significance of this correlation was evaluated using the cor.test function in R.

**Distance between eQTLs and transcription start site**

The distance between each significant eQTL and the transcription start site (TSS) of its associated eGene was calculated by comparing the variant position to the TSS position and strand of the gene according to Gencode v21. Positive distances indicate variants downstream from the TSS, negative distances indicate upstream variants. Comparison between groups of eQTLs was run using only the top eQTL per gene and the absolute value of distance from the associated gene's TSS.

**Overlap of eQTLs with H3K27ac**

We tested the global overlap between eQTLs and H3K27ac from human fetal, infant and adult dorsal frontal cortex and cerebellum and embryonic cortex from BrainSpan (Li et al., 2018) and Reilly et al (Reilly et al., 2015) . Intersection between sets of coordinates were performed using Bedtools (Quinlan and Hall, 2010). We tested three sets of variants: (1) best eQTLs per eGene,

(2) all significant eQTL with FDR< 0.05 in the corresponding eQTL category of "Prenatal", "Postnatal" or "Constant", and (3) a background group composed by all variants tested for eQTL, excluding those with a P<0.05 to any gene at any time period tested. 95% confidence intervals were obtained by bootstrapping variants 100 times (Figure S4).

**Enrichment of eQTLs in functional genomic elements**

We tested the enrichment of different categories of eQTLs in sets of genomics elements using GREGOR (Schmidt et al., 2015). We tested all best eQTL per gene in: (1) dorsal frontal cortex H3K27ac peaks from fetal and adult brain samples, and (2) 18 chromatin states whole-genome segmentation of sample E073-Medial Frontal Cortex Lobe from Epigenome Roadmap (Roadmap Epigenomics Consortium et al., 2015). We reported observed/expected number of overlapping eQTLs and BH adjusted P-values (Figure S4).

**Identification of rare protein-truncating variants**

Rare single nucleotide variants (SNVs), defined as not being observed in GnomAD (Lek et al., 2016) and only observed once in the 176 samples, were annotated using VEP with the LOFTEE plugin in Hail. Variants in genes that were not cortically expressed or that had zero reads in ≥10 samples in our data were excluded. Of the 128 variants marked as "stop gained", 110 were predicted to be high-confidence protein-truncating variants (PTVs) by LOFTEE, while 10 were in the last exon of the gene and predicted to be low-confidence PTVs (Table S5). Visual inspection of the PTV location within the protein-coding gene was used to define three categories of PTV (Nagy and Maquat, 1998; Rivas et al., 2015).

1. Probable NMD: A gene with multiple coding exons and the PTV is present in an exon included in all Gencode v21 isoforms and not in the last exon or with 50bp of the second to last exon.
2. Escaping NMD: A PTV in the last exon or fewer than 50bp from the splice site in the second to last coding exon.
3. Possible NMD: A PTV not meeting the criteria for the above two categories (i.e. a single exon gene or a PTV in an exon that differs between isoforms).

**Identification of rare deletions**

Rare copy-number deletions greater than 1kb size were identified as a subset of deletions predicted using MANTA version 1.3.1 with the default parameters. After removing deletions based on the non-PASS filter, there were a total of 98,522 deletions from 176 samples (560 deletions per sample). We defined rare deletions with following criteria: 1) Deletion never observed in 3,804 parents with whole-genome sequencing data (An et al., 2018); 2) Deletion only observed once in 176 samples. The 398 rare deletions were annotated by intersecting lists of transcript and exon annotations taken from the Gencode v21 database using the bedtools intersect module. After excluding non-coding deletions, genes that were not cortically expressed or that had zero reads in ≥10 samples in our data were excluded, 32 rare coding deletions remained. The raw coverage data and sequence reads were visually inspected to identify 20 high confidence deletions in cortically-expressed protein-coding genes (Table S5). The same criteria were used as for PTVs to estimate whether nonsense-mediated decay would be expected.

**Rare eQTL detection with allele specific expression**

For PTVs, the number of RNA-seq reads that contained the PTV was calculated using mpileup function in samtools. The percentage of reads with a variant was calculated as the number of reads with a PTV divided by the number of reads with a PTV or reference. The variant status could not be estimated for a small number of reads, which were excluded from the calculation. The percent of variant reads are shown in Table S5 and are plotted in Figure 5 of the main manuscript.

**Rare eQTL detection with using imputation of expected expression**

When a subject carries a PTV or deletion in one copy of an autosomal gene, it is generally assumed that the expression of that gene in the subject's tissue would be reduced by one half, relative to its expected value, due to nonsense mediated decay. To account for subject-specific variation in the measurement of gene expression we imputed the expected expression value, with the imputation based on expression values of genes found to be correlated from the population sample. To implement this approach, the 148 gene-subject pairs with a PTV or deletion (Table S5) were treated as a missing value and imputed by MICE imputation (van Buuren and Groothuis-Oudshoorn, 2011).

- Step 1: For each of 148 genes that have at least one missing values, we select 30 genes as predictors by conducting a Lasso regression on the non-missing genes using the lars R package; the proper tuning parameters are used so that 30 predictors are selected.
- Step 2: Estimate the conditional expressions of missing genes using the selected predictors with MICE R package. Get the expected expression by averaging over m imputation results (m=5 by algorithm default).
- Step 3: Compare the expected (imputed) expression E[x] with the measured true expressions x by calculating the difference d = E[x] - x, which corresponds to the $\log_2$ transformed ratio of expected expression to measure expression in the original natural scale. The differences in observed and expected expression are shown in Table S5 and are plotted in Figure 5 of the main manuscript.

The imputed expression values showed a similar range and distribution to that of the measured gene expression levels. The differences clustered around 0, indicating that the realized expression levels were largely similar to the expected levels.

**Gene sets associated with CNS traits and disorders**

To compare with 23,782 brain-expression genes, we created the list of gene sets for previous trait and disorder association and functional properties. Gene identifiers were converted between studies based on the complete HUGO Gene Nomenclature Committee dataset. Autism spectrum disorder (ASD) risk genes were obtained from Satterstrom et al. (Satterstrom et al., 2018), an exome sequencing based gene discovery refining to high-confidence genes (n=99) at a false discovery rate (FDR) ≤ 0.1. Genes associated with developmental delay (n=93) were selected from the exome sequencing analysis of the Deciphering Developmental Disorders project (Deciphering Developmental Disorders Study, 2017). From Heyne et al. (Heyne et al., 2018), we chose 33 genes as high-confidence epilepsy candidates, where multiple *de novo* variants were seen.

We used significant loci from the genome-wide association studies (GWAS) of attention deficit hyperactivity disorder (ADHD) (Demontis et al., 2019), Alzheimer's disease (Lambert et al.,

2013), educational attainment (EA) (Lee et al., 2018), schizophrenia (SCZ) (Schizophrenia Working Group of the Psychiatric Genomics Consortium, 2014), major depressive disorder (MDD) (Wray et al., 2018), multiple sclerosis (MS) (International Multiple Sclerosis Genetics Consortium et al., 2013), Parkinson's disease (PD) (Chang et al., 2017). We selected loci from a summary statistics file if publicly available, otherwise we used the table of genome-wide significant loci from each study. We retrieved genes harboring loci or within 10kb from loci. We excluded the extended MHC region on the chromosome 6, known to have a substantial number of genes due to high linkage disequilibrium (Schizophrenia Working Group of the Psychiatric Genomics Consortium, 2014) for downstream enrichment analyses.

Constrained genes were defined as probability of loss-of-function intolerant (pLI) score ≥ 0.995 in the ExAC database (Lek et al., 2016). Genes specific to cell types in the mid-fetal cortical development were selected from Nowakowski et al. (Nowakowski et al., 2017) and BrainSpan (Li et al., 2018). For all gene lists, see Table S6.

**Maximal period of expression in CNS trait and disorder associated gene lists**

To compare the expression of genes associated with CNS traits and disorders over time we defined the period of peak expression as the developmental period with the highest median $\log_2$CPM values across all periods for each gene. For each gene list the period of peak expression was annotated and the mean of this value across all genes was used to define the mean period of maximal expression. These values were used to define the order of traits and disorders (Figure 6A).

**Enrichment of DLPFC eQTLs in SNPs associated with complex phenotypes**

We tested for enrichment of DLPFC eQTLs among GWAS significant SNPs using a permutation-based procedure. GWAS SNPs were taken from published GWAS summary statistics at a significance threshold of $p < 5 \times 10^{-8}$. The following procedure was repeated separately for summary statistics from GWAS of four phenotypes: schizophrenia (Schizophrenia Working Group of the Psychiatric Genomics Consortium, 2014), educational attainment (Lee et al., 2018), multiple sclerosis (International Multiple Sclerosis Genetics Consortium et al., 2013), and Alzheimer's disease (Lambert et al., 2013). First, SNPs in the summary statistics and the list of SNPs tested for eQTL discovery were filtered to those in 1000 Genomes Project data. SNPs in the GWAS summary statistics were then filtered to variants also tested for eQTL discovery, using direct (same SNP) or proxy ($r^2$ >0.8 in CEU 1000 Genomes samples) as defined by the PLINK 1.9 – r2 command (Chang et al., 2015a). To estimate the proportion of eQTL SNPs among phenotype associated SNPs versus among all SNPs (or a sample of null SNPs), we recognized three important factors that could differ between null SNP sets and the phenotype associated SNPs: LD structure, MAF distribution, and gene density. First, to account for LD structure, we used PriorityPruner version 0.1.4 (http://prioritypruner.sourceforge.net) to LD prune SNPs supervised by GWAS p-value in order to preferentially retain as many phenotype-associated SNPs as possible while adequately removing SNPs in high LD ($r^2$ > 0.7 within a sliding 500kb window) using in 1000 Genomes CEU sample data. Second, the remaining SNPs were grouped according to their MAF decile. Third, each remaining SNP was grouped into decile of gene density to allow for differential opportunity to be identified as an eQTL. Gene density was determined by the number of genes within the 1MB eQTL detection window as defined by the annotation package "TxDb.Hsapiens.UCSC.hg38.knownGene" from R Bioconductor

(Bioconductor Core Team and Bioconductor Package Maintainer, 2016). SNPs in the ~3.2 MB HLA region (hg38 coordinates: chr6:29,751,784-32,915,731) as defined by the "GWASTools" R Bioconductor package (Gogarten et al., 2012) and UCSC genome browser (Kent et al., 2002) were excluded from enrichment testing. Next, 1 million null SNP sets were drawn by matching each phenotype associated SNP (GWAS $p < 5 \times 10^{-8}$) to a random SNP matched on both MAF and gene density. Enrichment fold statistics were computed as the proportion of eQTLs in the phenotype-associated set divided by the mean proportion of eQTLs across null sets. P-values were calculated as the proportion of null set fold-enrichment statistics as or more extreme than the observed phenotype-associated fold enrichment statistic. This permutation procedure was repeated for each of six eQTL SNP lists: all, general, prenatal-specific, prenatal non-specific, postnatal-specific, and postnatal non-specific.

**Gene-set analysis of eGenes and GWAS data**

To assess whether eQTL targets (eGenes) are enriched for GWAS signal, we performed competitive gene set enrichment analysis for each group of eGenes using the MAGMA software (de Leeuw et al., 2015). We input the eGene lists from prenatal, postnatal, general, non-specific prenatal, and non-specific postnatal (Table S4) and summary statistics from published GWAS of schizophrenia (Schizophrenia Working Group of the Psychiatric Genomics Consortium, 2014), autism spectrum disorder [doi: https://doi.org/10.1101/224774], educational attainment (Lee et al., 2018), multiple sclerosis (International Multiple Sclerosis Genetics Consortium et al., 2013), Alzheimers disease (Lambert et al., 2013), triglycerides (Willer et al., 2013), and height (Wood et al., 2014). First, SNPs from the GWAS summary statistics files were annotated to NCBI protein-coding genes that passed RNA-seq QC in our DLPFC expression data with a 10kb window on either side of the gene boundaries. Next, a gene-level analysis was done to determine the strength of association for each gene with phenotype of interest (equation here). To assess whether genes in the eGene gene-sets are more strongly associated with the phenotype of interest than other genes, gene-based z-scores are regressed on a gene-set indicator variable and MAGMA default covariates (gene size, gene density, sample size, 1/MAC, and the log of each of these) (equation here). The beta coefficient for the gene-set indicator variable is tested for significance $H_A: \beta_1 > 0$ . Results from this analysis are reported in Table S6. We did not observe significant enrichment of GWAS signal from any of the six phenotypes tested in any of the five temporally assigned eGene gene sets. However, we repeated a similar test, annotating GWAS summary statistics SNPs to NCBI protein-coding genes with 10kb flanking region that passed RNA-seq QC in our DLPFC expression data and had an assigned pLI score, and found that a gene-set defined as pLI score >= 0.995 showed significant enrichment for stronger GWAS association in all tested phenotypes except for multiple sclerosis, compared to genes with pLI score < 0.995.

**Co-localization analysis of CNS traits and disorders**

Coloc2 (https://github.com/Stahl-Lab-MSSM/coloc2) (Dobbyn et al., 2018), an updated version of coloc (Giambartolomei et al., 2014), was used to formally test for co-localization of GWAS signal from schizophrenia (Schizophrenia Working Group of the Psychiatric Genomics Consortium, 2014) and educational attainment (Lee et al., 2018) summary statistics with our five eQTL categories. Coloc2 tests five hypotheses (H0: no association, H1: GWAS association only, H2: eQTL association only, H3: both but not co-localized, H4: both and co-localized) and returns a posterior probability for each hypothesis in each region. Posterior probably of H4>= 0.8 is strong Bayesian evidence of co-localization.

### DATA AND SOFTWARE AVAILABILITY

Open source scripts used in this manuscript are referenced throughout. The pipeline for whole-genome analysis is available online at: https://github.com/sanderslab/psychcore-compute-platform. Raw RNA-seq and WGS data, along with processed files, are available at http://psychencode.org.

### Supplemental Figures

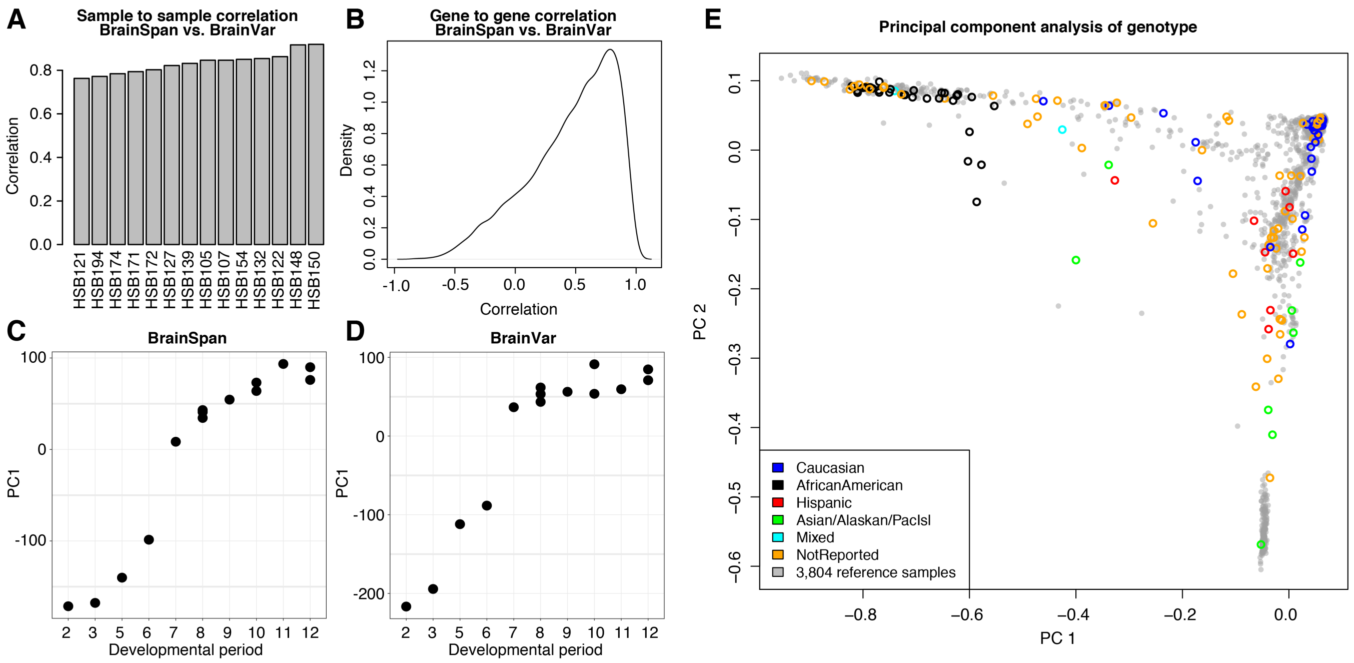


**Figure S1. Comparison of RNA-seq data between BrainSpan and BrainVar and WGS ancestry prediction. A)** Pearson correlation coefficients of gene expression for 14 overlapping samples with RNA-seq data generated from the dorsolateral frontal cortex in BrainSpan (Li et al., 2018) and BrainVar. **B)** Distribution of Pearson correlation coefficients between 23,782 cortically-expressed genes for RNA-seq data for 14 overlapping samples in BrainSpan (Li et al., 2018) and BrainVar. **C)** First principal component of gene expression by developmental period for all samples in the BrainSpan dataset (Li et al., 2018). **D)** First principal component of gene expression by developmental period for all samples in BrainVar. **E)** Principal component analysis using common variation called from WGS data for all 176 samples in BrainVar against a reference of 3,804 independent parents with WGS data from the Simons Simplex Collection (An et al., 2018). Self-reported ancestry is indicated by color for the BrainVar samples.

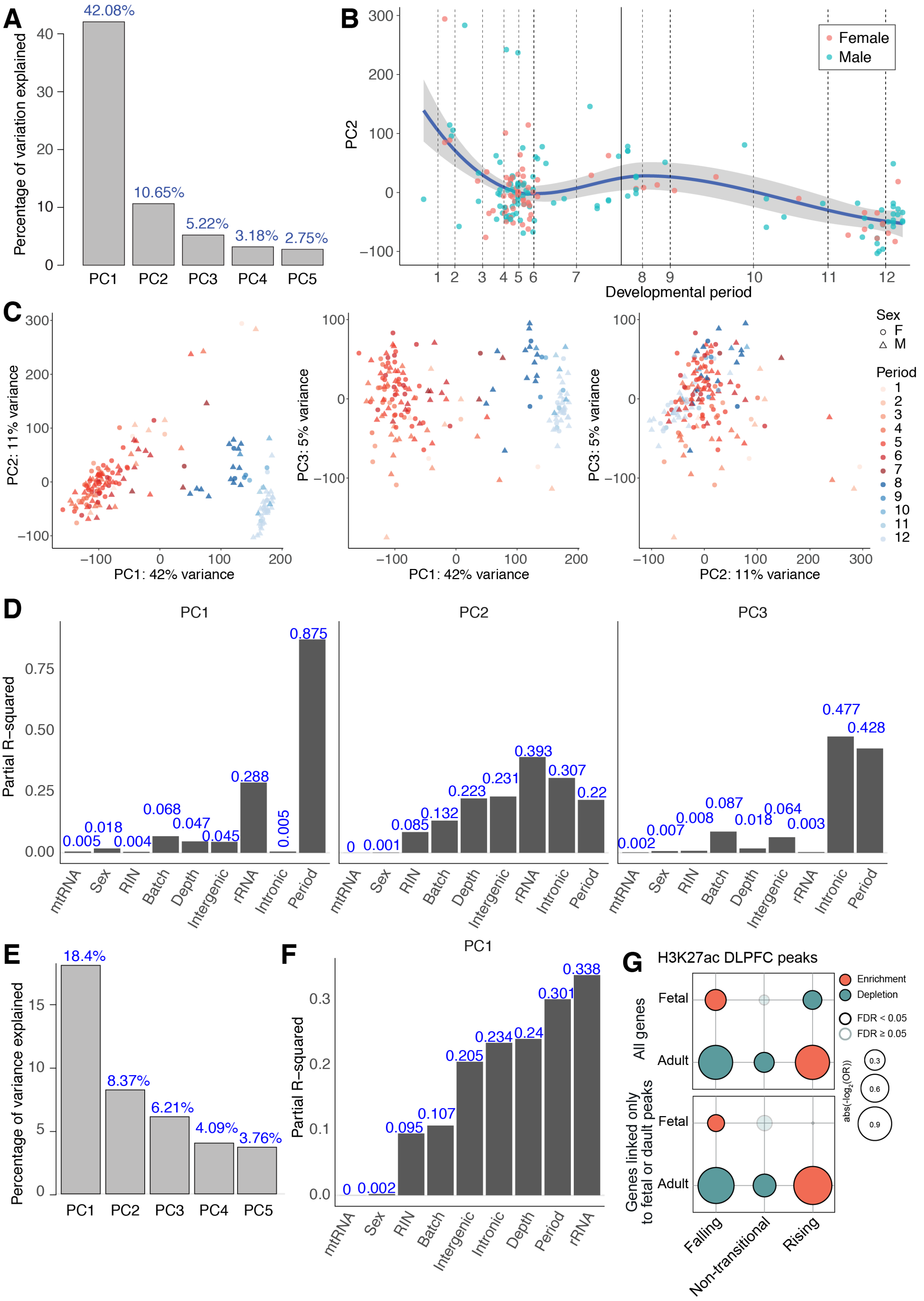
A
Percentage of variation explained
42.08%
10.65%
5.22%
3.18%
2.75%
PC1 PC2 PC3 PC4 PC5
B
PC2
Female
Male
Developmental period
1 2 3 4 5 6 7 8 9 10 11 12
C
PC2: 11% variance
PC1: 42% variance
PC3: 5% variance
PC1: 42% variance
PC3: 5% variance
PC2: 11% variance
Sex
F
M
Period
1 2 3 4 5 6 7 8 9 10 11 12
D
PC1
PC2
PC3
Partial R−squared
0.005 0.018 0.004 0.068 0.047 0.045 0.288 0.005 0.875
0 0.001 0.085 0.132 0.223 0.231 0.393 0.307 0.22
0.002 0.007 0.008 0.087 0.018 0.064 0.003 0.477 0.428
mtRNA Sex RIN Batch Depth Intergenic rRNA Intronic Period
E
Percentage of variance explained
18.4%
8.37%
6.21%
4.09%
3.76%
PC1 PC2 PC3 PC4 PC5
F
PC1
Partial R−squared
0 0.002 0.095 0.107 0.205 0.234 0.24 0.301 0.338
mtRNA Sex RIN Batch Intergenic Intronic Depth Period rRNA
G
H3K27ac DLPFC peaks
Enrichment
Depletion
FDR < 0.05
FDR ≥ 0.05
abs(−log$_2$(OR))
0.3
0.6
0.9
All genes
Genes linked only to fetal or dault peaks
Fetal
Adult
Falling
Non-transitional
Rising

**Figure S2. Principal component analysis of gene expression. A)** A scree plot showing the variance in gene expression explained by the first five principal components (PC1 to PC5) across all samples and 23,782 cortically-expressed genes in the BrainVar dataset. **B)** The relationship between PC2 (y-axis) and developmental period (x-axis); the equivalent plot for PC1 is shown in Figure 2A of the main manuscript. **C)** Relationship between PC1, PC2, and PC3 for each sample (points) with developmental period indicated by color and genotypic sex by symbol. **D)** Correlation between PC1, PC2, and PC3, known variables, and residuals. **E)** Trajectory analysis identified 12,077 genes involved in the late-fetal transition and 11,705 Non-transitional genes (Figure 2B). To assess the extent to which the late-fetal transition explains the temporal variance captured by PC1 in the initial analysis, we repeated the principal component analysis for the 11,705 Non-transitional genes. PC1 of this secondary analysis explains on 18.4% of the variance in gene expression. **F)** The correlation between PC1 and known variables and residuals for the secondary analysis based on Non-transitional genes only. **G)** The enrichment for H3K27ac peaks, detected in the fetal and adult human dorsolateral frontal cortex in BrainSpan (Li et al., 2018), with Falling, Non-transitional and Rising genes. The analysis is shown for all 23,782 cortically-expressed genes (top) and limited to genes with an associated H3K27ac peak during at least one developmental stage.

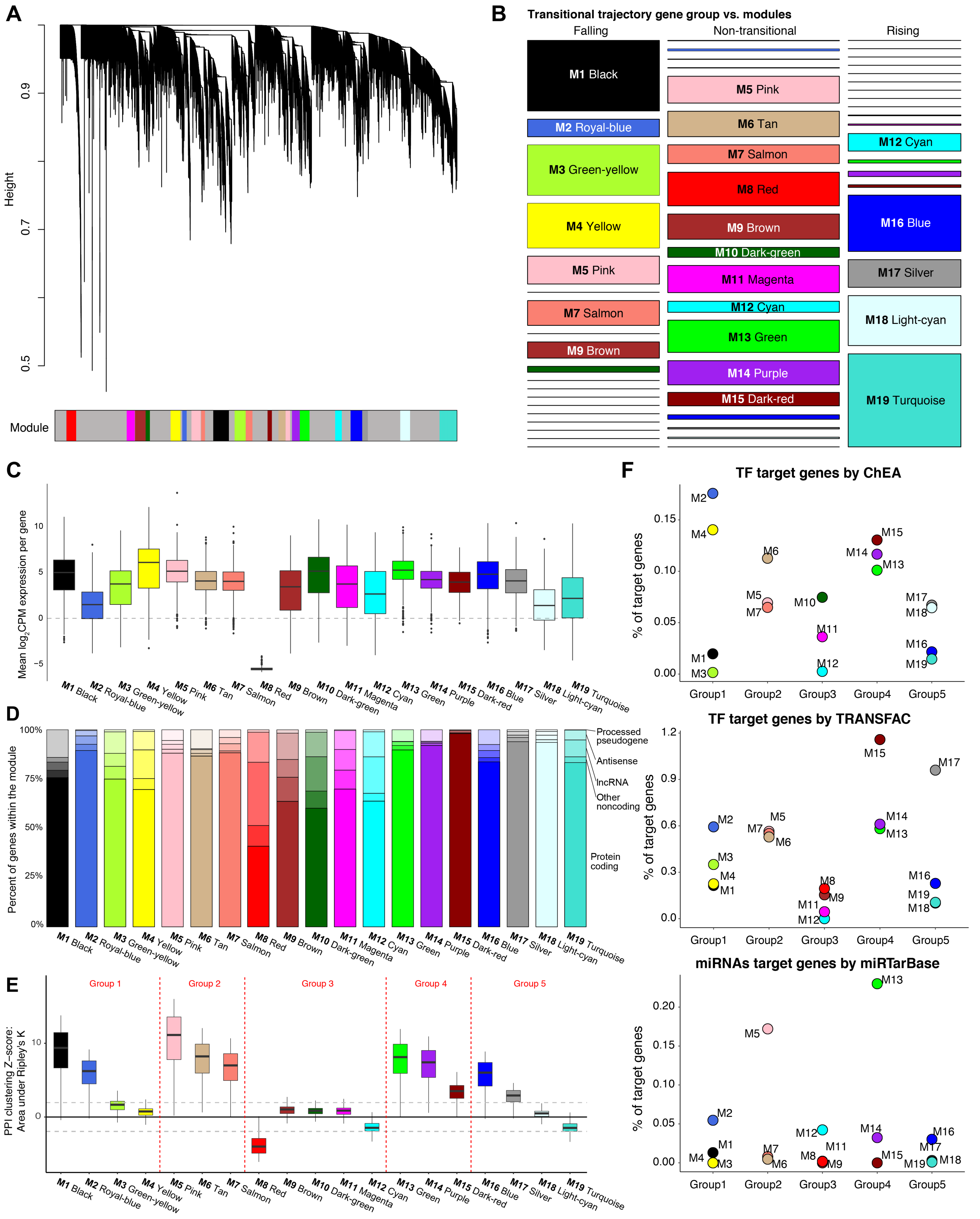
A
Height
Module
B
Transitional trajectory gene group vs. modules
Falling
Non-transitional
Rising
M1 Black
M2 Royal-blue
M3 Green-yellow
M4 Yellow
M5 Pink
M6 Tan
M7 Salmon
M8 Red
M9 Brown
M10 Dark-green
M11 Magenta
M12 Cyan
M13 Green
M14 Purple
M15 Dark-red
M16 Blue
M17 Silver
M18 Light-cyan
M19 Turquoise
C
Mean log2CPM expression per gene
D
Percent of genes within the module
Processed pseudogene
Antisense
lncRNA
Other noncoding
Protein coding
E
PPI clustering Z-score: Area under Ripley's K
Group 1
Group 2
Group 3
Group 4
Group 5
F
TF target genes by ChEA
TF target genes by TRANSFAC
miRNAs target genes by miRTarBase
% of target genes
Group1
Group2
Group3
Group4
Group5

**Figure S3. WGCNA module relationship and characteristics. A)** Dendrogram of 10,459 cortically-expressed genes assigned to 19 WGCNA modules; 13,323 genes were not assigned to any module (grey). **B)** A mosaic plot showing the overlap of genes between modules (y-axis and color) and the three transitional trajectory gene sets (x-axis), with the area of each rectangle representing the number of overlapping genes. **C)** Median $log_2$CPM expression of genes within each module (x-axis and color) across all 176 samples in the cohort. **D)** The percentage of protein-coding and noncoding genes (lncRNA, Antisense, Processed pseudogenes, and other noncoding shown by opacity) within each module. **E)** The connectivity within BioGRID protein-protein interaction (PPI) networks of the genes within each module is shown as a Z-score distribution (Cornish and Markowetz, 2014) by permutation against all 23,782 cortically-expressed genes. **F)** The percentage of genes in each module targeted by transcription factors (TFs) predicted by ChIP Enrichment Analysis (ChEA, (Lachmann et al., 2010), top) and TRANSFAC (http://genexplain.com/transfac/, middle) and the percentage of genes targeted by miRNA prediction from the mirTarbase database ((Chou et al., 2018), bottom).

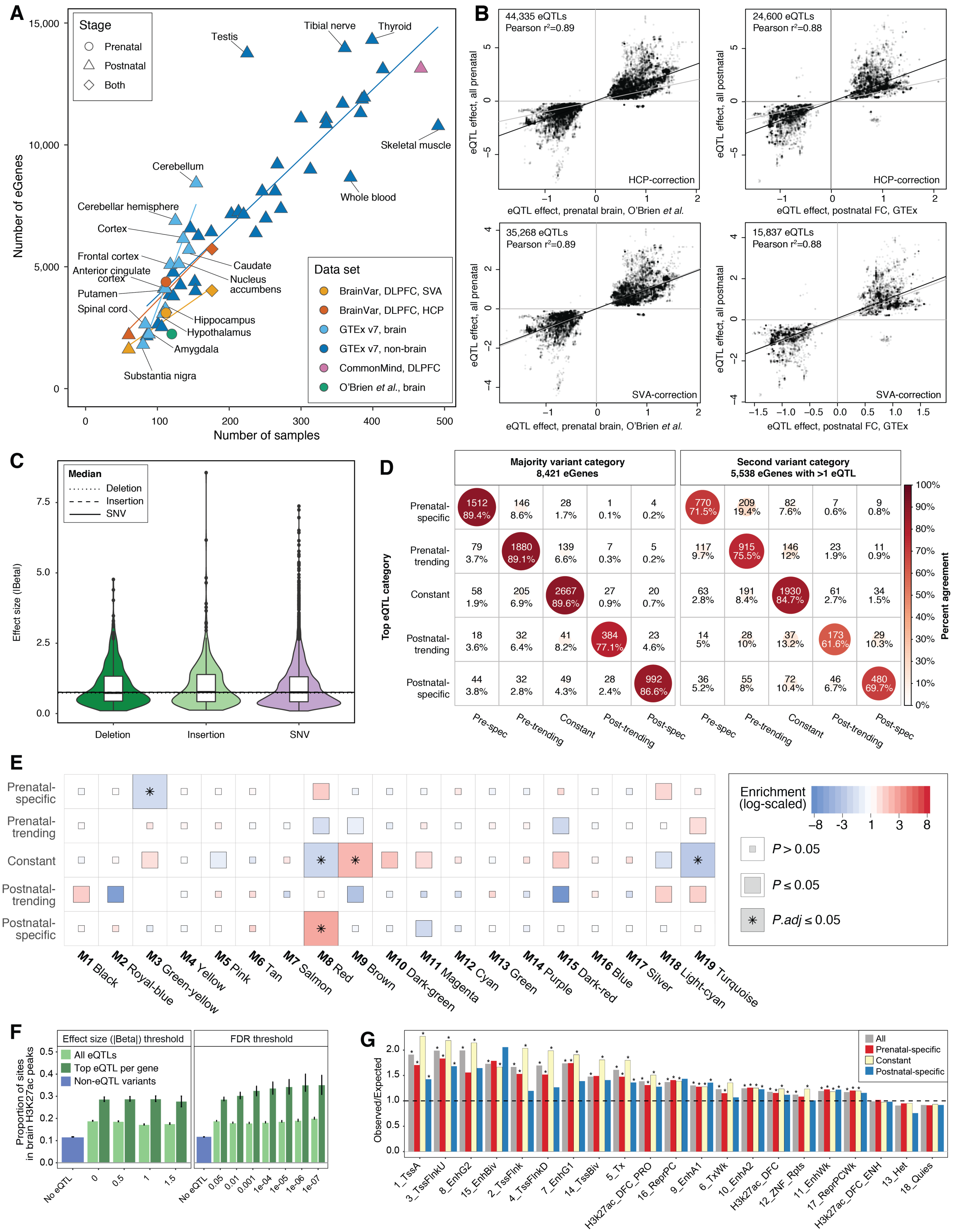
A
Stage
Prenatal
Postnatal
Both
Number of eGenes
Number of samples
Testis
Tibial nerve
Thyroid
Skeletal muscle
Whole blood
Cerebellum
Cerebellar hemisphere
Cortex
Frontal cortex
Anterior cingulate cortex
Putamen
Spinal cord
Caudate
Nucleus accumbens
Hippocampus
Hypothalamus
Amygdala
Substantia nigra
Data set
BrainVar, DLPFC, SVA
BrainVar, DLPFC, HCP
GTEx v7, brain
GTEx v7, non-brain
CommonMind, DLPFC
O'Brien et al., brain
B
44,335 eQTLs
Pearson r²=0.89
HCP-correction
eQTL effect, all prenatal
eQTL effect, prenatal brain, O'Brien et al.
24,600 eQTLs
Pearson r²=0.88
eQTL effect, all postnatal
eQTL effect, postnatal FC, GTEx
35,268 eQTLs
Pearson r²=0.89
SVA-correction
15,837 eQTLs
Pearson r²=0.88
C
Median
Deletion
Insertion
SNV
Effect size (|Beta|)
D
Majority variant category
8,421 eGenes
Second variant category
5,538 eGenes with >1 eQTL
Top eQTL category
Prenatal-specific
Prenatal-trending
Constant
Postnatal-trending
Postnatal-specific
Pre-spec
Pre-trending
Constant
Post-trending
Post-spec
Percent agreement
1512 89.4%
146 8.6%
28 1.7%
1 0.1%
4 0.2%
770 71.5%
209 19.4%
82 7.6%
7 0.6%
9 0.8%
79 3.7%
1880 89.1%
139 6.6%
7 0.3%
5 0.2%
117 9.7%
915 75.5%
146 12%
23 1.9%
11 0.9%
58 1.9%
205 6.9%
2667 89.6%
27 0.9%
20 0.7%
63 2.8%
191 8.4%
1930 84.7%
61 2.7%
34 1.5%
18 3.6%
32 6.4%
41 8.2%
384 77.1%
23 4.6%
14 5%
28 10%
37 13.2%
173 61.6%
29 10.3%
44 3.8%
32 2.8%
49 4.3%
28 2.4%
992 86.6%
36 5.2%
55 8%
72 10.4%
46 6.7%
480 69.7%
E
Prenatal-specific
Prenatal-trending
Constant
Postnatal-trending
Postnatal-specific
M1 Black
M2 Royal-blue
M3 Green-yellow
M4 Yellow
M5 Pink
M6 Tan
M7 Salmon
M8 Red
M9 Brown
M10 Dark-green
M11 Magenta
M12 Cyan
M13 Green
M14 Purple
M15 Dark-red
M16 Blue
M17 Silver
M18 Light-cyan
M19 Turquoise
Enrichment (log-scaled)
P > 0.05
P ≤ 0.05
P.adj ≤ 0.05
F
Effect size (|Beta|) threshold
FDR threshold
All eQTLs
Top eQTL per gene
Non-eQTL variants
Proportion of sites in brain H3K27ac peaks
No eQTL
G
All
Prenatal-specific
Constant
Postnatal-specific
Observed/Expected
1_TssA
3_TssFlnkU
8_EnhG2
15_EnhBiv
2_TssFlnk
4_TssFlnkD
7_EnhG1
14_TssBiv
5_Tx
H3K27ac_DFC_PRO
16_ReprPC
9_EnhA1
6_TxWk
10_EnhA2
H3K27ac_DFC
12_ZNF_Rpts
11_EnhWk
17_ReprPCWk
H3K27ac_DFC_ENH
13_Het
18_Quies

**Figure S4. eGene concordance and characteristics. A)** Scatterplot and best fit lines for the number of eGenes (y-axis) against sample size (x-axis) reported in BrainVar and several published data sets (Fromer et al., 2016; O'Brien et al., 2018; The GTEx Consortium et al., 2017). **B)** Scatterplot of the eQTL direction and magnitude of effect for variant-gene pairs reaching FDR-significance in BrainVar and published datasets (O'Brien et al., 2018; The GTEx Consortium et al., 2017). The plots on the left compare BrainVar to prenatal whole brain (O'Brien et al., 2018), while those on the right compare BrainVar to adult DLPFC (The GTEx Consortium et al., 2017). The plots at the top are for eQTLs discovered in the BrainVar prenatal samples, while those at the bottom are eQTLs discovered in BrainVar postnatal samples. Best fit line is plotted in black, slope of 1 is plotted in gray. **C)** Distribution of the absolute value of eQTL effect size (regression beta) for all eQTLs binned by deletions, insertions, and single nucleotide variants (SNV). **D)** The temporal specificity of an eGene was defined by the eQTL with the lowest p-value (top eQTL). To assess the consistency of this approach, we assessed the number of eGenes where the temporal specificity of the top eQTL matched the majority of eQTLs for the eGene (left) or the eQTL with the second lower p-value (right). **E)** Enrichment of 19 WGCNA modules with eGenes, divided into five categories based on temporal specificity. **F)** Proportion of eQTLs that overlap an H3K27ac peak detected in human DLPFC in BrainSpan (Li et al., 2018). Results are binned by eQTL effect size (left) and FDR threshold (right) showing all eQTLs and the top eQTL per gene (shown by shade). **G)** Barplot of observed versus expected overlap using GREGOR analysis between temporally specific and Constant eQTLs with functional loci from 18 chromatin states defined by the Roadmap Epigenome Project (https://egg2.wustl.edu/roadmap/web_portal/chr_state_learning.html#exp_18state).

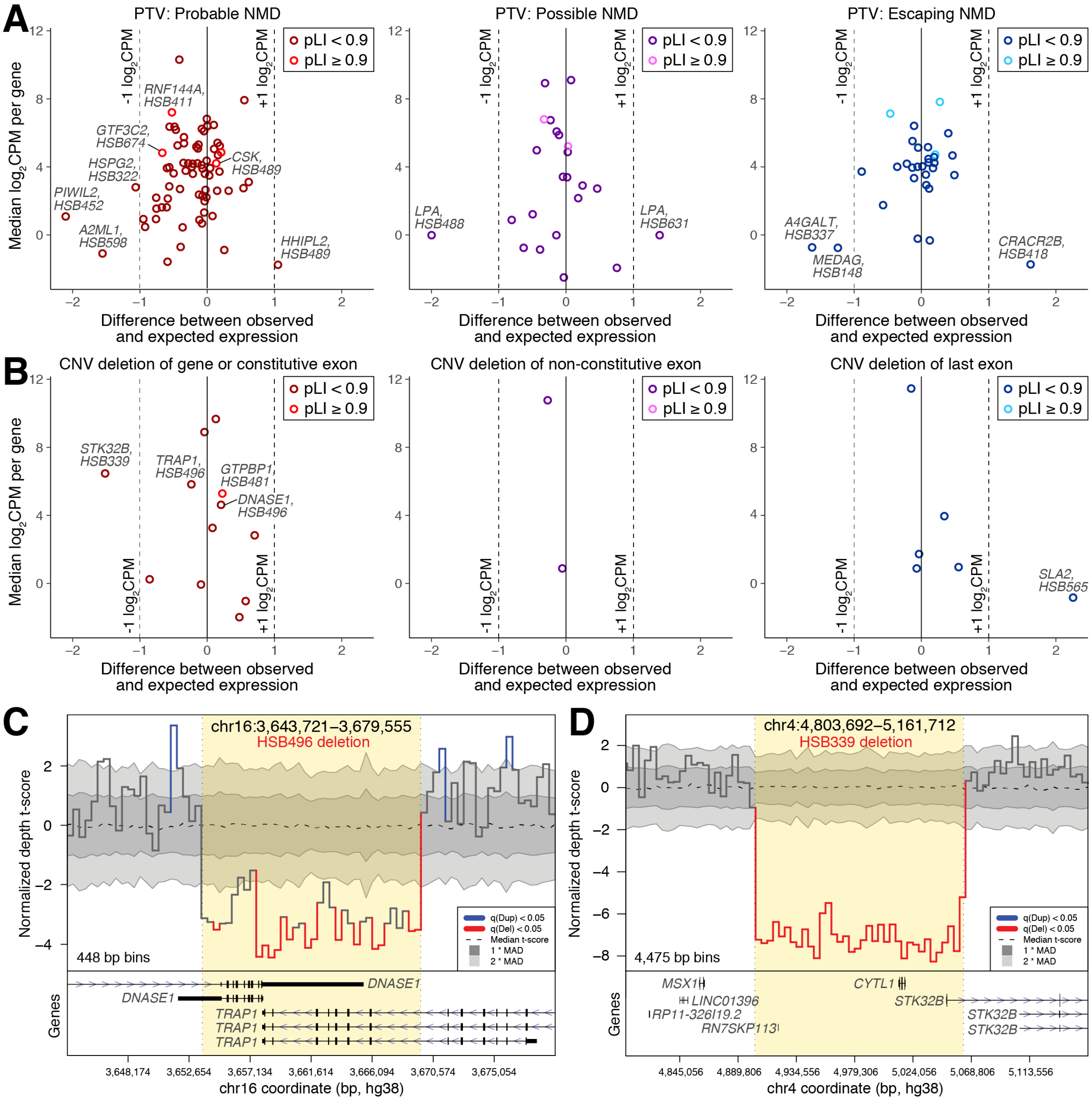


**Figure S5. Rare variant eQTLs. A)** A) The difference in observed vs. expected expression (x-axis) against the median $log_2CPM$ expression of the gene in all samples (y-axis) is shown for 74 gene-sample pairs in which one sample has a PTV expected to cause nonsense mediated decay, "Probable NMD" (left), 23 gene-sample pairs with “Possible NMD” PTVs in single exon genes or with the PTV-containing exon varying between isoforms (middle), and for 31 gene-sample pairs with “Escaping NMD” PTVs in the last exon or 50bp upstream of the last splice site (right). Selected genes and samples are labeled. **B)** Corresponding plots are shown for rare deletions affecting one or more constitutive exons (12 genes in 11 deletions, left), nonconstitutive exons (2 deletions, middle), or the last exon (6 deletions, right). **C)** The depth of coverage in the WGS is shown for sample HSB496 compared to the coverage of the other samples. A region deleted is indicated by yellow shading and the genes in the region are shown at the bottom. The effect of this CNV on expression is shown in Figure 5B of the main manuscript. **D)** A similar plot of a deletion in sample HSB339, gene STK32B, which is markedly decreased in expression (panel 'B').

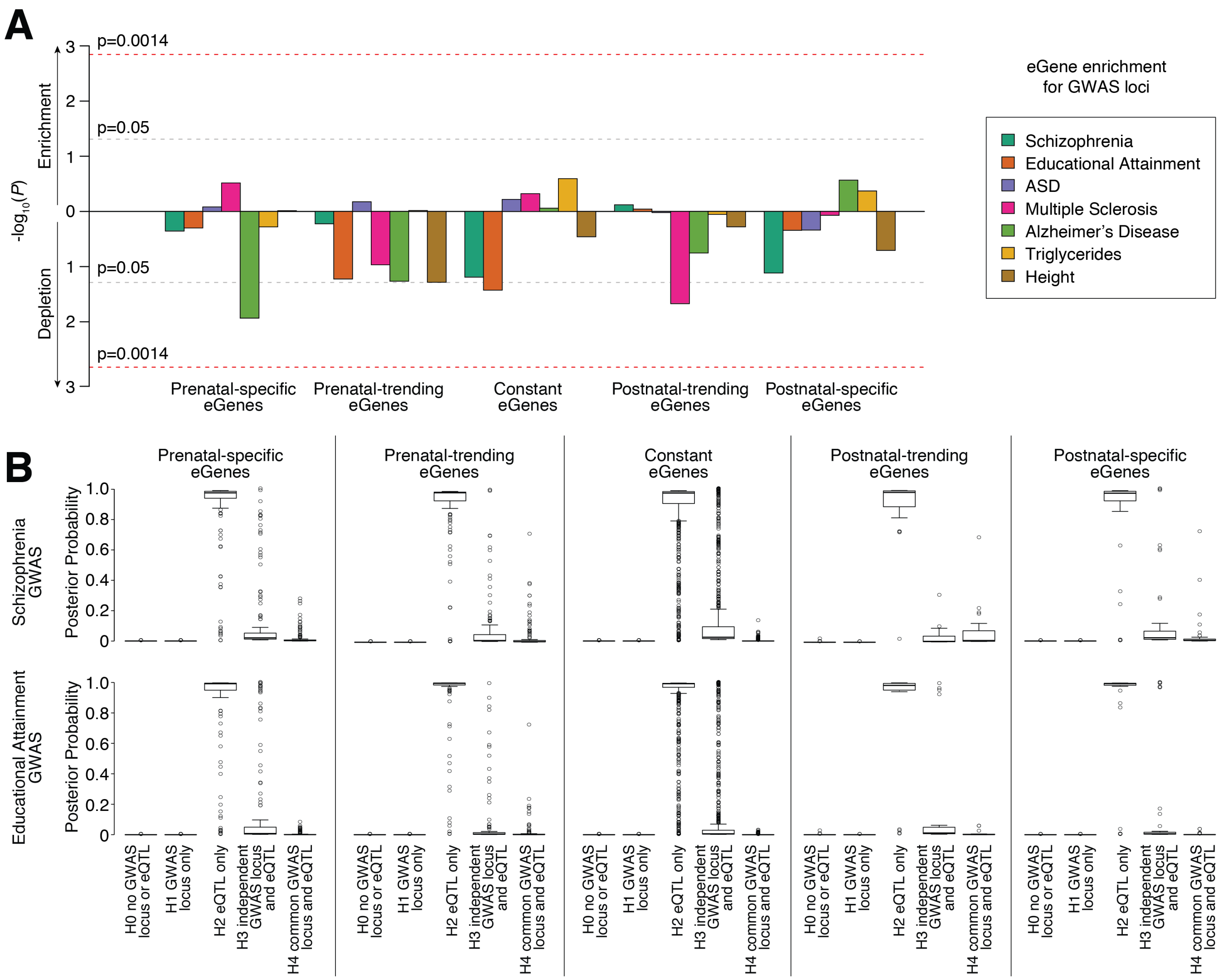


**Figure S6. eGene enrichment and eQTL co-localization with complex traits and disorders. A)** Competitive gene-set enrichment analysis testing for enrichment of GWAS signal among categories of eGenes, correcting for linkage disequilibrium. **B)** Posterior probabilities from tests of co-localization between GWAS and eQTL signal for schizophrenia and educational attainment. Statistical analysis: A: Multi-marker analysis of genomic annotation (MAGMA) B: Bayesian test for co-localization using summary statistics (Coloc2).

### Supplemental Tables

**Table S1.** Sample information. (Also see Figure 1, S1)

**Table S2.** Gene information. (Also see Figures 2, 3, 4, 6, 7, S2, S3, S4, S6, S7)

**Table S3.** WGCNA results. (Also see Figure 3, S3)

**Table S4.** Common cis-eQTL results. (Also see Figure 4, S4)

**Table S5.** Rare variants eQTLs. (Also see Figure 5, S5)

**Table S6.** Gene set enrichment for CNS trait and disorder genes. (Also see Figure 6)

**Table S7.** GWAS-eQTL enrichment. (Also see Figure 7)
